# Dietary fibre reverses adverse post-stroke outcomes in mice via short-chain fatty acids and its sensing receptors GPR41, GPR43 and GPR109A

**DOI:** 10.1101/2023.05.15.540735

**Authors:** Alex Peh, Evany Dinakis, Hamdi Jama, Dovile Anderson, Darren J. Creek, Gang Zheng, Michael de Veer, Charles R. Mackay, Tenghao Zheng, Barbara K. Kemp-Harper, Brad R.S. Broughton, Francine Z. Marques

**Affiliations:** Hypertension Research Laboratory, School of Biological Sciences, Monash University, Melbourne, Australia; Cardiovascular & Pulmonary Pharmacology Group, Department of Pharmacology, Monash University, Melbourne, Australia; Monash Institute of Pharmaceutical Sciences, Monash University, Melbourne, Australia; Monash Biomedical Imaging, Monash University, Clayton, Victoria, Australia; Department of Microbiology, Biomedical Discovery Institute, Faculty of Medicine, Nursing and Health, Monash University, Clayton, Victoria, Australia; School of Pharmaceutical Sciences, Shandong Analysis and Test Center, Qilu University of Technology (Shandong Academy of Sciences), Jinan, 250014, China; Heart Failure Research Group, Baker Heart and Diabetes Institute, Melbourne, Australia; Victorian Heart Institute, Monash University, Melbourne, Australia

**Author notes:** Contributed equally as senior authors (Brad R.S. Broughton, Francine Z. Marques). **Correspondence to:** A/Prof Francine Marques, Hypertension Research Laboratory, School of Biological Sciences, Faculty of Science, Monash University, Melbourne, Australia. E.

**Keywords:** Ischaemic stroke, gut microbiome, short-chain fatty acids, microbiota, metabolites

## Abstract

Dietary fibre intake is associated with fewer cases of ischaemic stroke. This is likely via the microbiota-gut-brain axis, where fibre is fermented by the gut microbiota, releasing short-chain fatty acids (SCFAs). However, whether fibre or SCFAs can reverse adverse post-stroke outcomes remains unknown. Here, we demonstrated that a low fibre diet exacerbates post-stroke outcomes in mice. This was reversed by a high fibre diet or direct supplementation with SCFAs (delivered either in the water or a high SCFA-releasing diet) immediately after stroke. These modulated the gut microbiome and improved the gut epithelial barrier integrity, which was associated with fewer activated neutrophils and more neuroblast cells in the brain. We then investigated the SCFA-receptors GPR41/43/109A using a triple knockout mouse model, which exhibited poorer stroke outcomes and recovery. These results show that post-stroke interventions using dietary fibre and/or SCFA supplementation, acting via GPR41/43/109A signalling, may represent new therapeutic strategies for stroke-induced brain injury.

## Introduction

More than 12 million people suffer a stroke each year, with approximately 50% of patients resulting in death ^1^. Current treatments, such as recombinant tissue plasminogen activator and endovascular clot retrieval, must be administered within a limited time window (t-PA: typically, 4.5 hours and endovascular clot retrieval: 24 hours), which restricts the number of patients that are eligible to ∼10%. Furthermore, both treatments only target blood clot removal and do not target cell death or repair mechanisms to improve post-stroke outcomes. Therefore, there is an urgent need to identify new therapeutic strategies for stroke.

Experimental and clinical studies over the past decade have provided overwhelming evidence that the microbiota-gut-brain axis significantly impacts stroke outcomes^2^. Stroke is often associated with intestinal dysmotility, gut microbiota dysbiosis, and disruption of the intestinal epithelial barrier, which results in poor stroke prognosis^3, 4^. One of the significant roles of the gut microbiota is to harvest available micronutrients and energy for the host by fermenting dietary fermentable fibre, such as some types of soluble fibre and resistant starches, which reach the large intestine intact^5^. This fermentation results in the release of microbial metabolites known as short-chain fatty acids (SCFAs), such as acetate, butyrate, and propionate, as by-products.

SCFAs have anti-inflammatory properties^6^ and support the maintenance of the gut epithelial barrier^7, 8^. A Westernised diet low in fibre leads to the proliferation of bacteria that digest the intestinal mucus layer, leading to the breakdown of the gut epithelial barrier^9^. Moreover, low fibre intake may irreversibly reduce the diversity of the gut microbiota and ultimately lead to the loss of specific beneficial bacterial species in the gut^10^. These changes may lead to intestinal dysfunction, increasing the development of chronic inflammation and the risk of stroke^11^. Evidence supports that increased dietary fibre intake or healthy gut microbiota lowers the risk of stroke^12, 13^. A meta-analysis found that dietary fibre intake was significantly associated with reduced risk of primary stroke occurrence^14^. A randomised clinical trial using an acetate- and butyrate-enriched high amylose maize starch (HAMSAB) was recently shown to lower systolic blood pressure^15^, a key risk factor for stroke, thus potentially reducing stroke risk. However, there is no evidence to date to suggest whether a high fibre diet or SCFAs supplementation may improve poor post-stroke recovery caused by a sustained low fibre diet prior to the onset of the stroke event.

One of the mechanisms of action of SCFAs is via the G-protein-coupled receptors (GPCRs) GPR41, GPR43 and GPR109A^16^. Activation of these receptors that are commonly found in the gut and immune cells^16^ contributes to the maintenance of the gut epithelial barrier, increases mucus production, and prevents inflammation^6, 17^. The interaction between SCFAs and their receptors on intestinal endocrine cells promotes the transmission of indirect signals to the brain by inducing intestinal hormone secretion through the vagal nerve pathway or into the bloodstream^18^. In addition, single-gene knockout mouse models of GPR43 and GPR109A show increased inflammation^19, 20^. Thus, it is possible that treatment with SCFAs could improve stroke-induced gut epithelial barrier dysfunction and reduce inflammation via these receptors.

In this study, we aimed to demonstrate the importance of dietary fermentable fibre in stroke prevention and recovery, and to investigate whether dietary fibre or supplementation with SCFAs improves poor post-stroke outcomes. Moreover, we aimed to determine whether post-stroke recovery depends on the SCFAs receptors GPR41, GPR43 and GPR109A, and some of the associated mechanisms. Importantly, our data demonstrate that low fibre intake leads to a worse stroke outcome. This can be reversed by high fibre/SCFA supplementation, acting via GPR41/43/109A, having an important translational potential as a new post-stroke intervention.

## Results

### High fibre-fed mice had better post-stroke outcomes

To determine the impact of fibre on stroke-induced outcomes, we fed mice control or nutrient-matched diets containing low or high fibre for 28 days before photothrombotic (PT) stroke surgery and for 7 days after the surgery (Fig. 1a). At day one (D1) post-stroke, high fibre-fed mice had a smaller cerebral infarct volume (Fig. 1b,c) and cerebral oedema (Fig. 1d) compared to low fibre-fed mice. Brain infarct volume (Fig. 1e,f) and oedema (Fig. 1g) remained significantly smaller in mice on a high fibre diet compared to mice on a low fibre diet at day seven (D7) post-stroke. To determine the infarct recovery rate from D1 to D7, the percent change in infarct volume was calculated. These findings were consistent with high fibre-feeding resulting in a greater infarct (Fig. 1h) and reduced oedema recovery rate (Fig. 1i) 7-days after PT stroke.

**Fig. 1:**
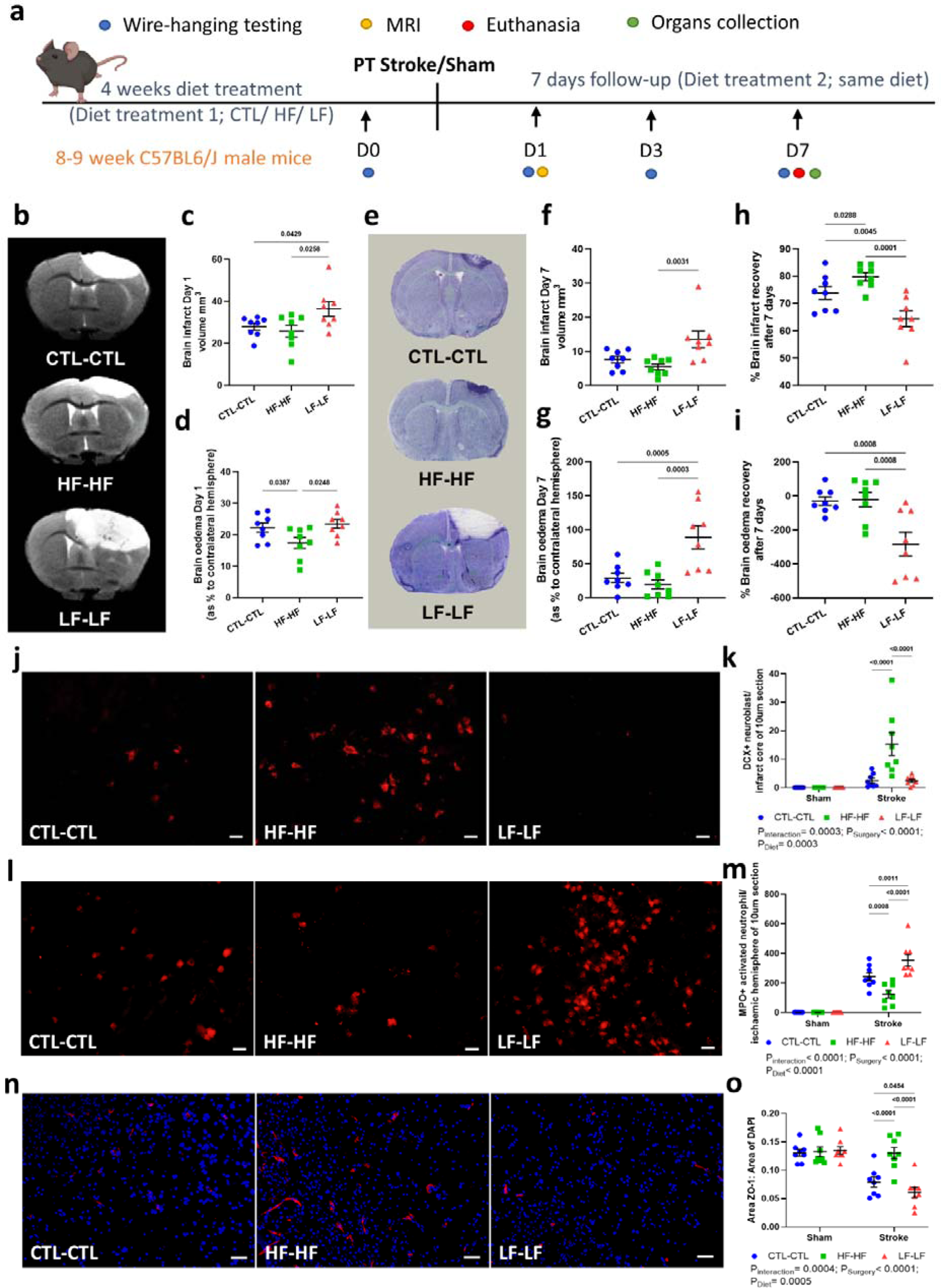
Dietary fibre is beneficial for stroke prevention and post-stroke recovery. **a**, The experimental design of the study. **b**, Representative images of brain infarct size at D1 (MRI-scanned brain: infarct region is the white oedema mass and unaffected areas are grey). Quantification of **c**, brain infarct volume and **d**, brain oedema at D1 post-stroke. **e**, Representative images of brain infarct size at D7 (thionin-stained brain: infarct region was shown in the white and unaffected area in purple) post-stroke. Quantification of **f**, brain infarct volume and **g**, brain oedema at D7 post-stroke. The recovery rate for **h**, brain infarct volume and **i**, brain oedema at D7 were calculated. Sample size= 8/group; error bar denotes mean±SEM. One-way ANOVA with FDR-adjusted was used to analyse data. **j**, Representative images of DCX+ neuroblast (scale bar=20µm) and **k**, the quantification of DCX+ neuroblast in the infarct core of CTL-CTL, HF-HF and LF-LF mice (n=8 mice per group; two-way ANOVA with FDR-adjusted). **l**, Representative images of MPO+ activated neutrophils (scale bar=20µm) and **m**, the quantification of MPO+ activated neutrophils in the infarct core of CTL-CTL, HF-HF and LF-LF mice (n=8 mice per group; two-way ANOVA with FDR-adjusted). **n**, Representative images of ZO-1 tight junction protein (red: ZO-1; blue: DAPI, scale bar=50µm) and **o**, the quantification of ZO-1 tight junction protein expression in the infarct core of CTL-CTL, HF-HF and LF-LF mice (n=8 mice per group; two-way ANOVA with FDR-adjusted). Error bar denotes mean±SEM. *Legend: CTL, control; HF, high fibre; LF, low fibre.

Although no difference in the latency to fall was found in sham mice performing the wire-hanging functional test (Extended Data Fig. 1), all post-stroke mice exhibited a significant impairment in their ability to hold onto the wire at D1 post-stroke (Extended Data Fig. 1). At D3 and D7 post-stroke, high fibre-fed mice had improved grip strength as they held onto the wire significantly longer than mice fed a low fibre diet (Extended Data Fig. 1).

We observed a significant increase in DCX+ neuroblasts, a marker of neurogenesis, in the infarct region of high fibre-fed mice compared to control and low fibre-fed mice at D7 following stroke (Fig. 1j,k). Moreover, we saw a significant reduction of MPO+ activated neutrophils in the infarct core of mice on a high fibre diet compared to mice on a control and low fibre diet (Fig. 1l,m). High fibre-fed mice also exhibited a significant increase in zonulim-1 (ZO-1) tight junction protein expression in the brain (Fig. 1n,o). However, no difference in activated neutrophils, neuroblast cells and tight junction protein expression was observed in the sham group, irrespective of the diet treatment (data not shown). In addition, we found that mice fed a high fibre diet resulted in ZO-1 tight junction protein expression remaining similar to high fibre-fed sham mice, however but significantly higher than control and low fibre post-stroke (Fig. 2a,b). Irrespective of their surgery groups, a high fibre diet resulted in longer villi (Fig. 2c-f) and fewer goblet cells (Fig. 2g-i), as well as longer colon and small intestine lengths (Fig. 2j,k). However, we found no difference in muscularis propria layer thickness, fibrosis thickness and various organs weights, including spleen, heart, liver, lung, and kidney (Extended Data Fig. 2).

**Fig. 2:**
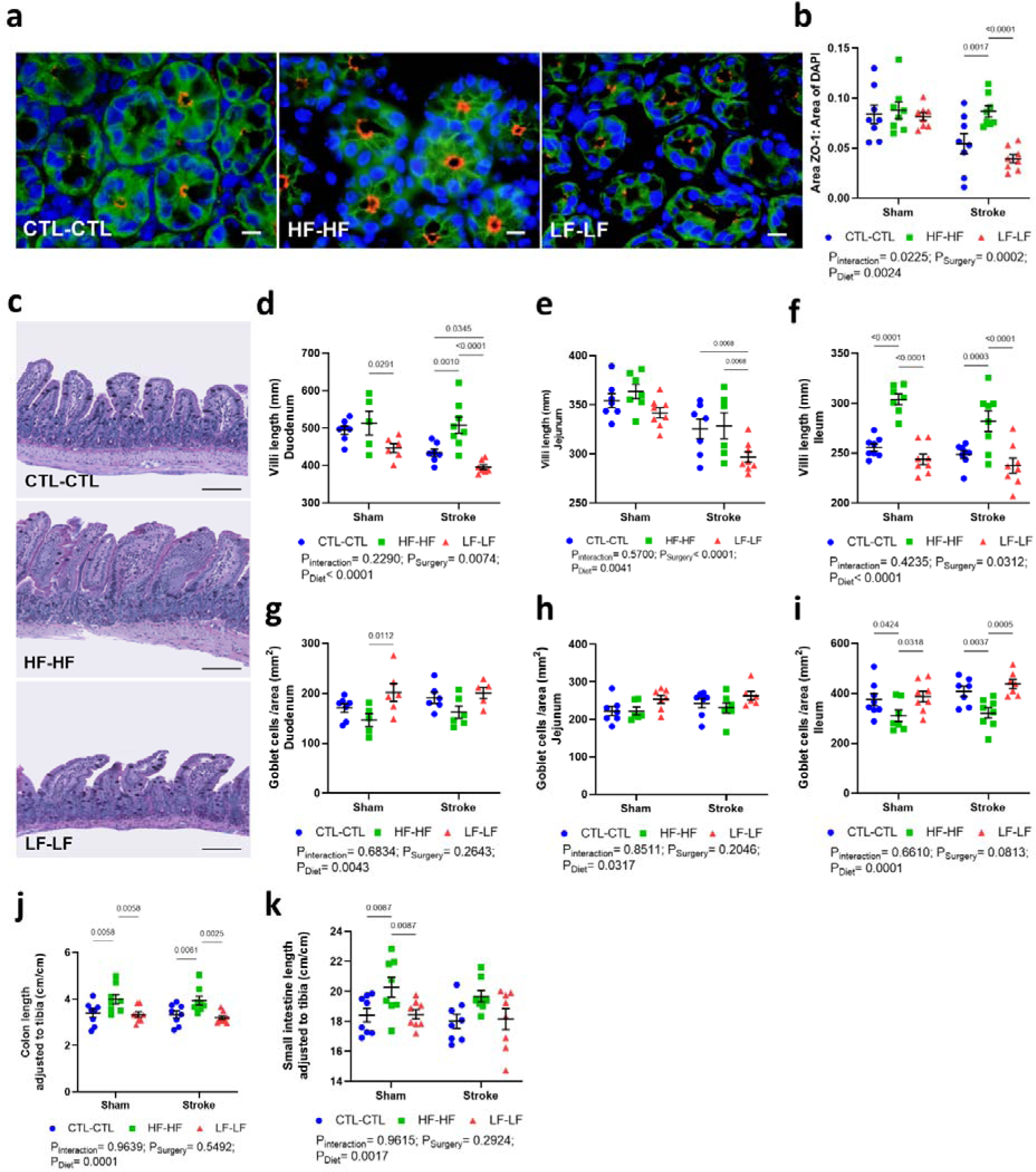
High fibre improved gut structure. **a**, Representative images of ZO-1 labelling in colon tissue of CTL-CTL, HF-HF and LF-LF mice post-stroke. Red: ZO-1; Green: EpCam (CD326); Blue: DAPI. Scale bar=10µm. **b**, Quantification of ZO-1 expression on the colon tissue of CTL-CTL, HF-HF and LF-LF mice post-surgery (n=8 mice per group). **c**, Representative images of staining of the ileum region showing the changes in villi length and goblet cells post-stroke stained using Periodic Acid-Schiff/Alcian blue. Scale bar: 100µm. Quantification of villi length in **d**, duodenum, **e**, jejunum, and **f**, ileum. Quantification of goblet cell number in **g**, duodenum, **h**, jejunum, and **i**, ileum. **j**, The length of the colon and **k**, the small intestine (SI) normalised to tibia length (cm/cm). Sample size= 8/group; error bar denotes mean±SEM. Two-way ANOVA was used to analyse data. *Legend: CTL, control; HF, high fibre; LF, low fibre.

To identify the changes in the gut microbiome post-stroke, 16S rRNA sequencing was performed. The sampling depth was captured at 8,000 reads, and the α-rarefaction curve indicated sufficient sequencing depth across all samples (data not shown). High fibre mice had the lowest metrics for α-diversity (Extended Data Fig. 3a-d), including Faith’s phylogenetic diversity, Shannon, observed species and species evenness index. The principal coordinate analysis (PCoA) using both weighted (Extended Data Fig. 3e) and unweighted UniFrac distances (Extended Data Fig. 3f) showed conspicuous clustering of gut bacterial communities based on their diet but not surgery type, showing that diet had a more profound impact on the microbiome than the surgery. In addition, phylum-level taxa plot distribution showed that the high fibre group had significantly more bacteria from phylum Bacteroidota (previously known as Bacteroidetes, q= 1.0542E-5) and fewer from Bacillota (previously known as Firmicutes, q= 2.7297E-5) (Extended Data Fig. 3g), irrespective of surgery group. When we further explored the abundance of species between diets using a linear discriminant analysis, we found that the genus *Bacteroides* was significantly more abundant in the high fibre group, while *Blautia*, *Bilophila*, *Enterococcus*, and *Lachnoclostridium* species were more abundant in the low fibre group (Extended Data Fig. 3h,i).

**Fig. 3:**
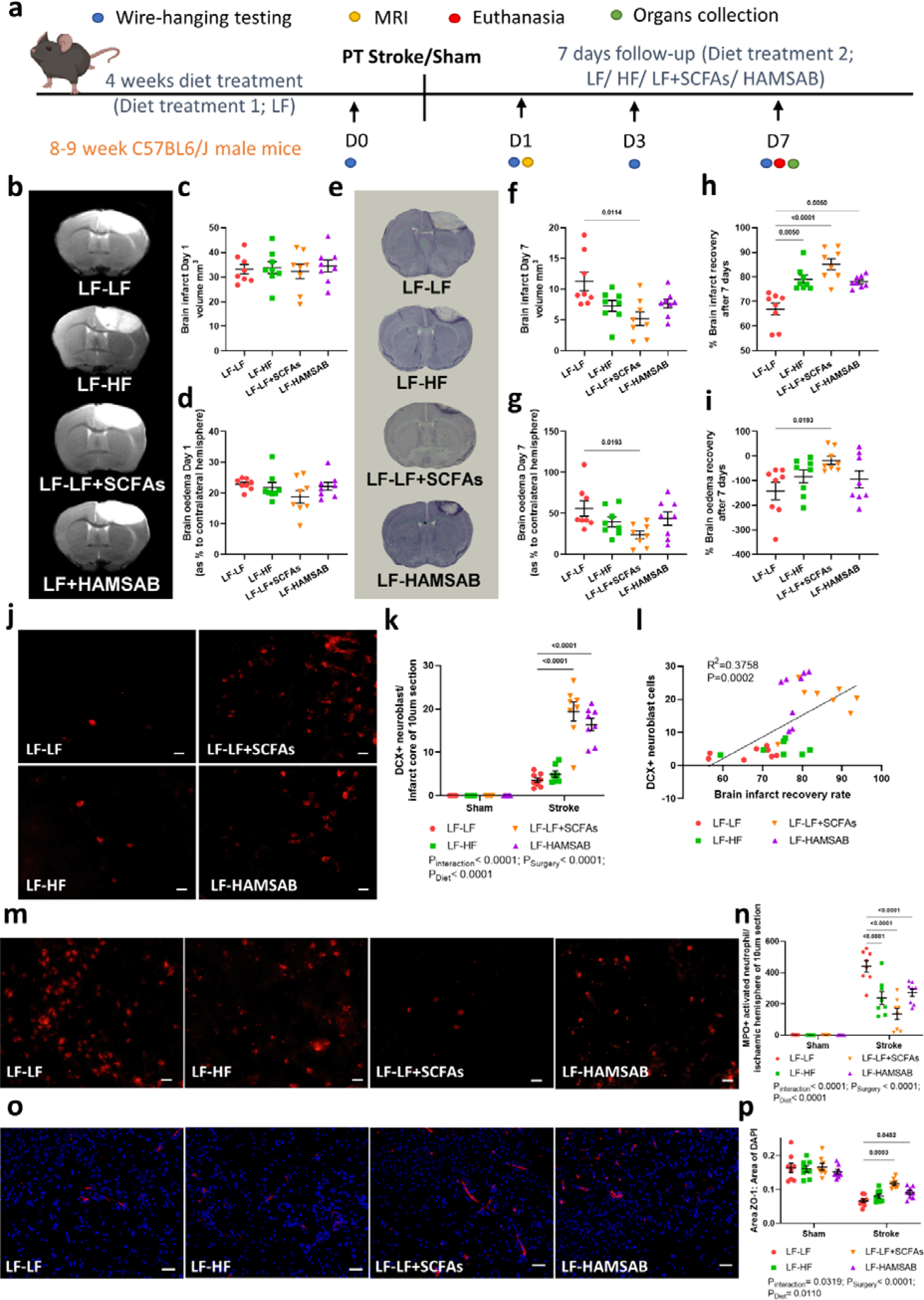
Gut microbial-produced metabolites reversed poor stroke outcomes. **a**, The experimental design of the study. **b**, Representative images of brain infarct size at D1 (MRI-scanned brain: infarct region is white and unaffected area grey). Quantification of **c**, brain infarct volume and **d**, brain oedema at D1 post-stroke. **e**, Representative images of brain infarct size at D7 (thionin-stained brain: infarct region was shown in the white and unaffected area in purple) post-stroke. Quantification of **f**, brain infarct volume and **g**, brain oedema at D7 post-stroke. The recovery rate for **h**, brain infarct volume and **i**, brain oedema at D7 were calculated. Sample size= 8/group; error bar denotes mean±SEM. One-way ANOVA with FDR-adjusted was used to analyse data. **j**, Representative images of DCX+ neuroblast (scale bar=20µm) and **k**, the quantification of DCX+ neuroblast in the infarct core of LF-LF, LF-HF, LF-LF+SCFAs and LF-HAMSAB mice (n=8 mice per group; two-way ANOVA with FDR-adjusted). **l**, Positive correlation between DCX+ neuroblast and brain infarct recovery rate. **m**, Representative images of MPO+ activated neutrophils (scale bar=20µm) and **n**, the quantification of MPO+ activated neutrophils in the infarct core of LF-LF, LF-HF, LF-LF+SCFAs and LF-HAMSAB mice (n=8 mice per group; two-way ANOVA with FDR-adjusted). **o**, Representative images of ZO-1 tight junction protein (red: ZO-1; blue: DAPI, scale bar=50µm) and **p**, the quantification of ZO-1 tight junction protein expression in the infarct core of LF-LF, LF-HF, LF-LF+SCFAs and LF-HAMSAB mice (n=8 mice per group; two-way ANOVA with FDR-adjusted). Error bar denotes mean±SEM. *Legend: LF, low fibre; HF, high fibre; SCFAs, short-chain fatty acids; HAMSAB, acetylated and butyrylated high amylose maise starch.

### Gut microbial-produced metabolites reversed poor stroke outcomes

Next, we aimed to determine whether supplementation with high fibre or SCFAs after ischaemic stroke could reverse the functional and anatomical impairment caused by extended low fibre feeding. Thus, mice were fed a low fibre diet for 4-weeks and then immediately post-surgery switched their diet to either a high fibre diet, a low fibre diet supplemented with SCFAs in the drinking water or a HAMSAB diet for 7-days (Fig. 3a). No difference in infarct volume (Fig. 3b,c) or brain oedema (Fig. 3d) was found across all groups at D1 post-stroke, thus, confirming a similar level of infarct damage occurred regardless of treatment. At D7, only SCFA-fed mice showed a significant reduction in infarct volume (Fig. 3e,f) and cerebral oedema (Fig. 3g). However, while high fibre, SCFAs and HAMSAB-fed groups had improved cerebral infarct recovery compared to low fibre mice (Fig. 3h), only SCFAs in the drinking water reduced brain oedema recovery (Fig. 3i).

In the wire-hanging test, there was no difference in the latency to fall times between diets in sham mice (Extended Data Fig. 4). However, high fibre, SCFAs and HAMSAB-fed stroke mice showed 4-6-fold improvement over the 7-days compared to low fibre stroke mice (Extended Data Fig. 4).

We found an increased number of neuroblasts in the cerebral infarct core of mice treated with SCFAs or fed a HAMSAB diet post-stroke compared to mice on a low fibre diet (Fig. 3j,k). Moreover, there was a positive correlation between the number of neuroblasts and brain infarct recovery (Fig. 3l). Furthermore, high fibre, SCFAs, and HAMSAB-fed mice showed significantly reduced neutrophil infiltration in the ischaemic hemisphere on D7 post-stroke (Fig. 3m,n). Additionally, SCFAs or a HAMSAB diet after PT stroke resulted in a significant increase in the expression of tight junctions in the brain compared to low fibre mice (Fig. 3o,p).

In the gut, ZO-1 expression in the colon was increased in mice fed a high fibre or HAMSAB diet as well as mice supplemented with SCFAs, suggesting that all three treatments improve gut epithelial integrity after stroke (Fig. 4a,b). These groups also had longer villi (Fig. 4c-f), fewer goblet cells (Fig. 4g-i), and a longer colon (Fig. 4j) and small intestine (Fig. 4k) than mice fed a low fibre post-stroke. However, we found no difference between groups in muscularis propria layer thickness, fibrosis thickness and liver weight. While we observed some difference in organ weight with treatment post-stroke, high fibre diet was the only group that prevented stroke-induced decrease in lung weight (Extended Data Fig. 5).

**Fig. 4:**
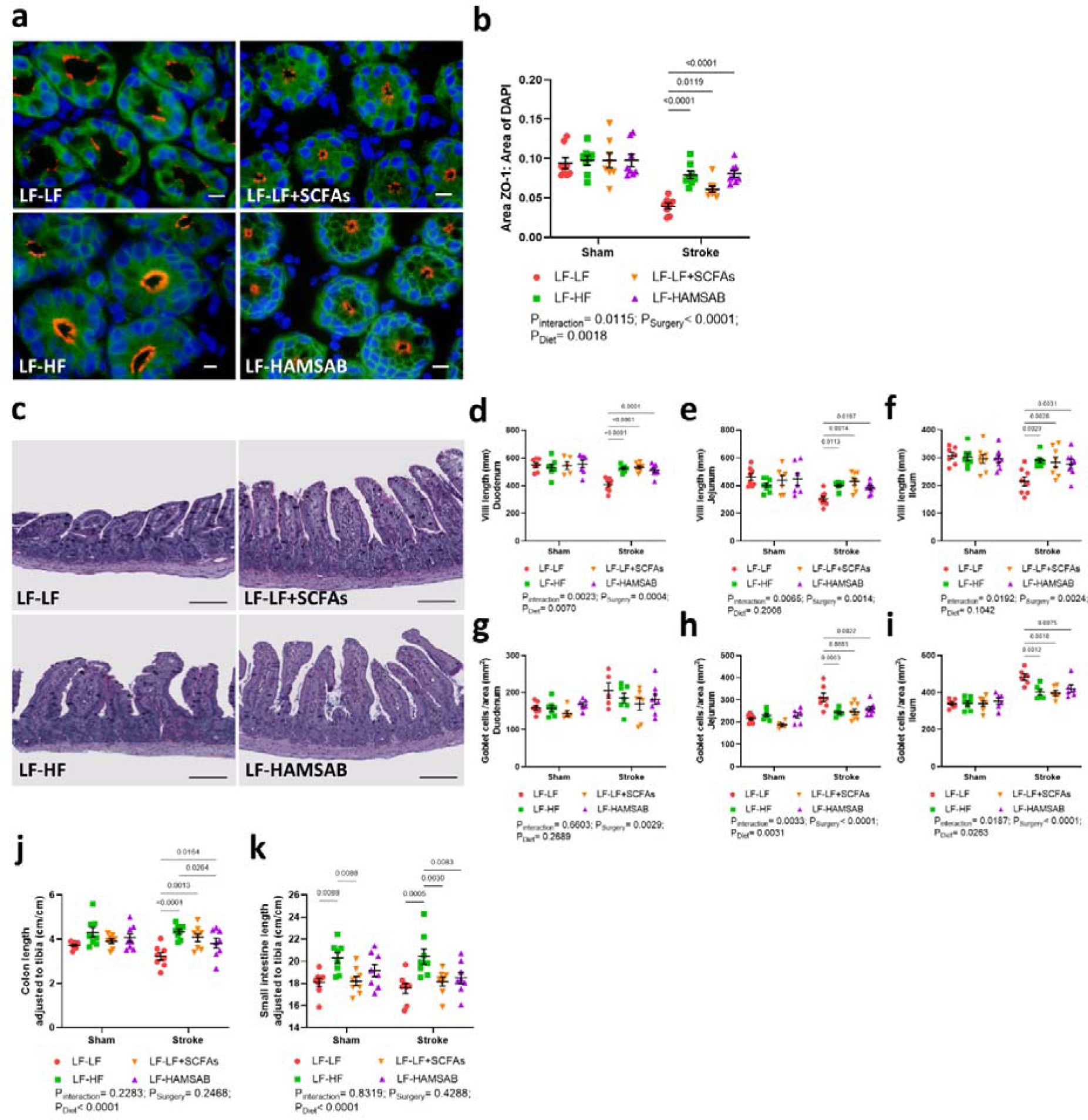
Gut microbial-produced metabolites improved gut structure. **a**, Representative images of ZO-1 labelling in colon tissue of LF-LF, LF-HF, LF-LF+SCFAs and LF-HAMSAB mice post-stroke. Red: ZO-1; Green: EpCam (CD326); Blue: DAPI. Scale bar=10µm. **b**, Quantification of ZO-1 expression on the colon tissue of LF-LF, LF-HF, LF-LF+SCFAs and LF-HAMSAB mice post-surgery (n=8 mice per group). **c**, Representative images of staining of the ileum region showing the changes in villi length and goblet cells post-stroke stained using Periodic Acid-Schiff/Alcian blue. Scale bar: 100µm. Quantification of villi length in **d**, duodenum, **e**, jejunum, and **f**, ileum. Quantification of goblet cell number in **g**, duodenum, **h**, jejunum, and **i**, ileum. **j**, The length of the colon and **k**, the small intestine (SI) normalised to tibia length (cm/cm). Sample size= 8/group; error bar denotes mean±SEM. Two-way ANOVA was used to analyse data. *Legend: LF, low fibre; HF, high fibre; SCFAs, short-chain fatty acids; HAMSAB, acetylated and butyrylated high amylose maise starch.

Similar to the results from previous feeding, α-diversity was higher for low fibre compared to the other groups (Extended Data Fig. 6a-d). We also found that post-surgery diet had an important effect on the gut microbiome, with clear clustering based on the type of post-surgery diet in both the weighted (Extended Data Fig. 6e) and unweighted UniFrac PCoAs (Extended Data Fig. 6f). At the phylum level, we found that the low fibre group had more Bacillota (q= 1.6074E-4) and Actinomycetota (q=4.7753E-5), while the SCFA group had more Bacteroidota (q= 1.6074E-4), Verrucomicrobia (q= 1.8583E-6) and Proteobacteria (q= 0.0016144), and the HAMSAB group had more Deferribacteres (q= 4.0041E-5) (Extended Data Fig. 6g). Additionally, more *Parabacteroides*, *Parasutterella*, *Akkermansia*, *Clostridioides* and *Enterococcus* were found in the SCFA group and more SCFA-producing bacteria such as *Faecalibaculum* and *Bacteroides* in the high fibre group (Extended Data Fig. 6h,i). However, we did not observe a change in SCFA levels in the circulation (Extended Data Fig. 7). Together, these findings demonstrated that high fibre and interventions with SCFAs can improve post-stroke outcomes caused by a pre-stroke low fibre diet.

### GPR41/43/109A knockout mice had worse stroke outcomes and recovery

To confirm the receptors by which dietary fibre, and subsequently SCFAs, mediate neuro-recovery post-stroke, GPR41/43/109A^-/-^ and wild-type (C57BL6/J) mice were fed a high fibre diet and subjected to PT stroke (Fig. 5a). GPR41/43/109A^-/-^ mice had a significantly larger infarct volume and brain oedema at D1 (Fig. 5b-d) and D7 (Fig. 5e-g) compared to C57BL6/J mice. Additionally, their infarct and oedema recovery were poor (Fig. 5h,i). Whilst there were no differences in the latency to fall times in all sham mice (Extended Data Fig. 8), times were significantly worse in GPR41/43/109A^-/-^ mice when compared to C57BL6/J mice post-stroke (Extended Data Fig. 8).

**Fig. 5:**
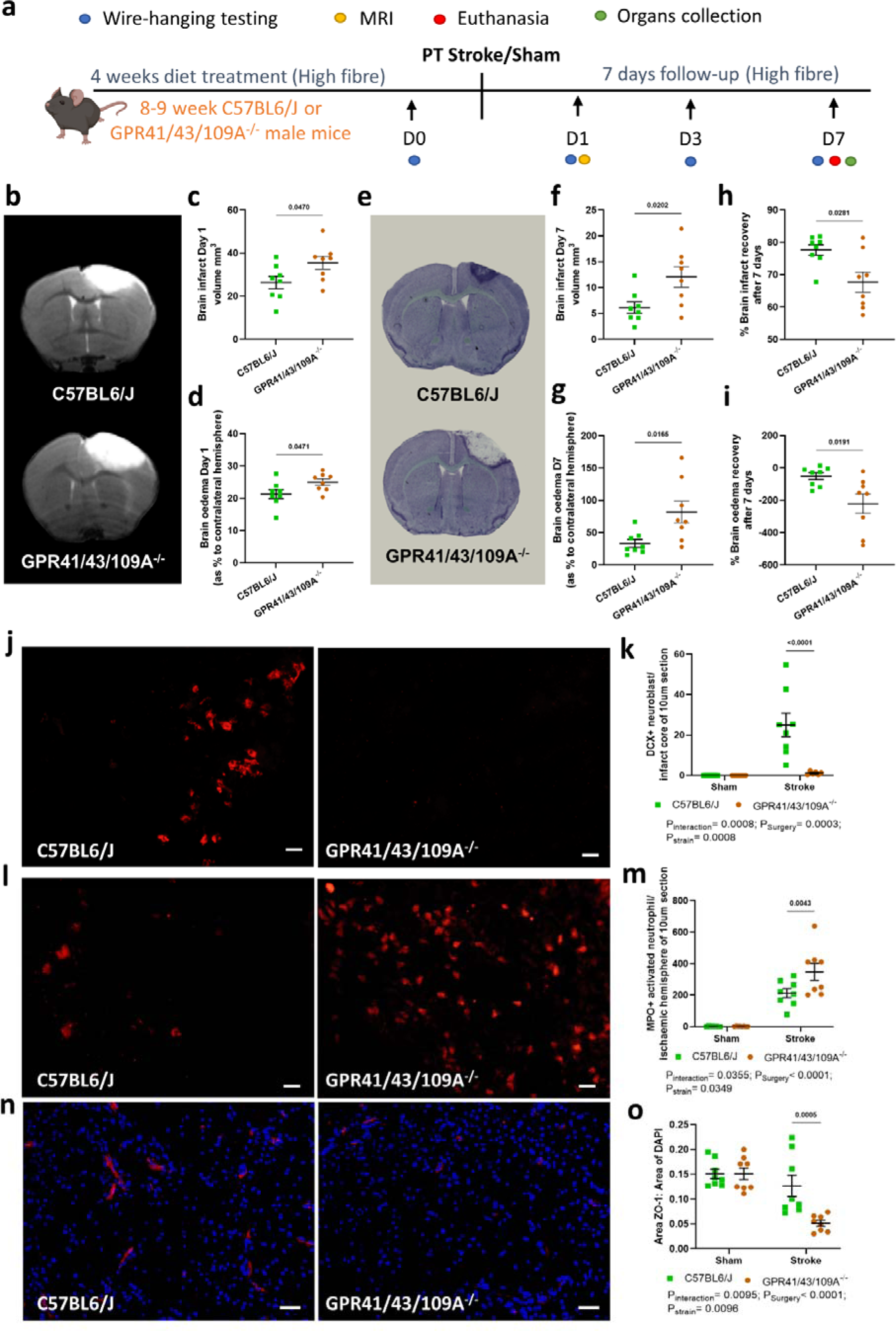
GPR41/43/109A knockout mice had worse stroke outcomes and recovery. **a**, The experimental design of the study. **b**, Representative images of brain infarct size at D1 (MRI-scanned brain: infarct region is white and unaffected area is grey). Quantification of **c**, brain infarct volume and **d**, brain oedema at D1 post-stroke. **e**, Representative images of brain infarct size at D7 (thionin-stained brain: infarct region was shown in the white and unaffected area in purple) post-stroke. Quantification of **f**, brain infarct volume and **g**, brain oedema at D7 post-stroke. The recovery rate for **h**, brain infarct volume and **i**, brain oedema at D7 were calculated. Sample size= 8/group; error bar denotes mean±SEM. One-way ANOVA with FDR-adjusted was used to analyse data. **j**, Representative images of DCX+ neuroblast (scale bar=20µm) and **k**, the quantification of DCX+ neuroblast in the infarct core of C57BL6/J and GPR41/43/109A^-/-^ mice (n=8 mice per group; two-way ANOVA with FDR-adjusted). **l**, Representative images of MPO+ activated neutrophils (scale bar=20µm) and **m**, the quantification of MPO+ activated neutrophils in the infarct core of C57BL6/J and GPR41/43/109A^-/-^ mice (n=8 mice per group; two-way ANOVA with FDR-adjusted). **n**, Representative images of ZO-1 tight junction protein (red: ZO-1; blue: DAPI, scale bar=50µm) and **o**, the quantification of ZO-1 tight junction protein expression in the infarct core of C57BL6/J and GPR41/43/109A^-/-^ mice (n=8 mice per group; two-way ANOVA with FDR-adjusted). Error bar denotes mean±SEM.

Using DCX as a marker for brain neurogenesis, we found few to no neuroblasts in GPR41/43/109A^-/-^ mice compared to C57BL6/J mice, indicating that these receptors are essential for fibre-induced neurogenesis (Fig. 5j,k). In addition, we observed a significant increase in MPO+ activated neutrophils in the ischaemic brain of GPR41/43/109A^-/-^ mice compared to C57BL6/J mice (Fig. 5l,m). Using the tight junction protein ZO-1 marker, we found little ZO-1 immunoreactivity in the brain of GPR41/43/109A^-/-^ mice compared to C57BL6/J mice (Fig. 5n,o). Moreover, ZO-1 expression in the colon was lower in the triple knockout mice (Fig. 6a,b). This suggests that gut epithelial barrier integrity is reduced in the absence of these receptors, even when the mice were fed a high fibre diet. Compared to C57BL6/J mice, GPR41/43/109A^-/-^ mice had shorter villi (Fig. 6c-f), more goblet cells (Fig. 6g-i), and shorter colon (Fig. 6j). However, we observed no difference in the small intestine length between GPR41/43/109A^-/-^ and C57BL6/J mice (Fig. 6k), as well as muscularis propria thickness, fibrosis thickness and various peripheral organ weights (Extended Data Fig. 9).

**Fig. 6:**
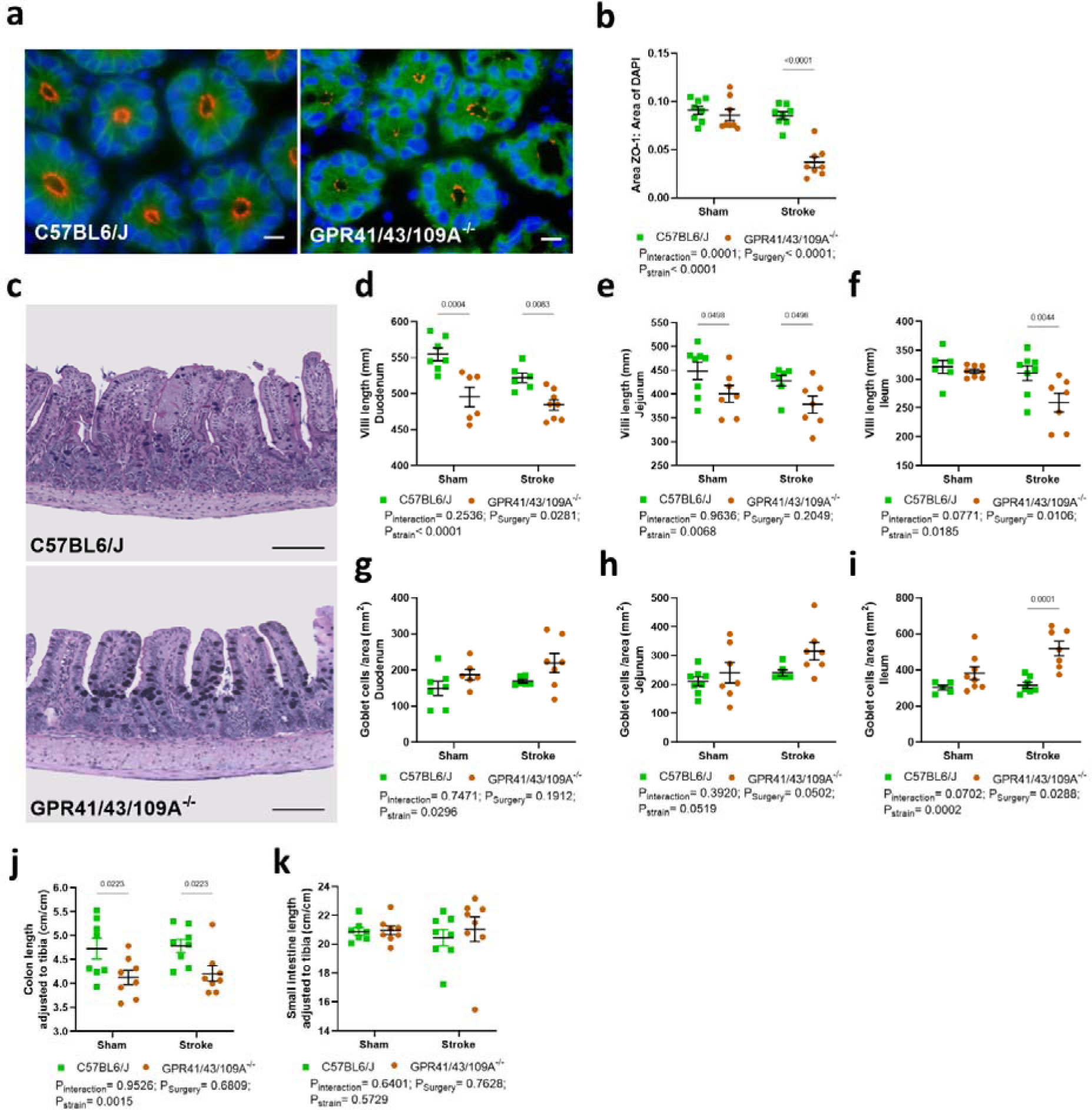
G protein-coupled receptors are essential to improve gut structure. **a**, Representative images of ZO-1 labelling in colon tissue of C57BL6/J and GPR41/43/109A^-/-^ mice post-stroke. Red: ZO-1; Green: EpCam (CD326); Blue: DAPI. Scale bar=10µm. **b**, Quantification of ZO-1 expression on the colon tissue of C57BL6/J and GPR41/43/109A^-/-^ mice post-surgery (n=8 mice per group). **c**, Representative images of staining of the ileum region showing the changes in villi length and goblet cells post-stroke stained using Periodic Acid-Schiff/Alcian blue. Scale bar: 100µm. Quantification of villi length in **d**, duodenum, **e**, jejunum, and **f**, ileumQuantification of goblet cell number in **g**, duodenum, **h**, jejunum, and **i**, ileum. The length of the **j**, colon and **k**, the small intestine (SI) normalised to tibia length (cm/cm). Sample size= 8/group; error bar denotes mean±SEM. Two-way ANOVA was used to analyse data.

Although α-diversity of the gut microbiome was similar between GPR41/43/109A^-/-^ and C57BL6/J mice (Extended Data Fig. 10a-d), the samples were clustered based on their strain in both weighted (Extended Data Fig. 10e) and unweighted UniFrac PCoAs (Extended Data Fig. 10f). However, we did not detect differences in microbial composition between the strains or the surgery (Extended Data Fig. 10g,h).

## Discussion

Dietary fibre is essential for nourishing the commensal bacteria in the large intestine, which produce beneficial metabolites such as SCFAs. These metabolites can be absorbed into the bloodstream and affect the host’s physiology by maintaining the gut epithelial barrier and reducing systemic inflammation^6, 21^. Here, we demonstrated that high fibre intake pre- and post-stroke improved recovery by restoring the integrity of the gut epithelial barrier, suppressing brain inflammation, and promoting neurogenesis. Conversely, mice with a sustained low fibre intake prior to stroke had a worse outcome and slower recovery. A previous study demonstrated that pre-stroke intake of SCFAs improved behavioural outcomes and modulated cortical network plasticity in mice^22^. However, it was unclear if poor stroke outcomes could be reversed with fibre and SCFAs. Nevertheless, we showed that post-stroke intake of high fermentable fibre and/or SCFAs (delivered either in the water or using a high SCFA-releasing diet) reversed adverse stroke outcomes within a short-term period, even if the pre-stroke diet was low in fibre. Finally, using a new triple knockout mouse model lacking all three major SCFA-sensing receptors, we demonstrate that this stroke-protective effect is dependent on GPR41/43/109A. These findings have important implications for post-stroke recovery and could lead to new, easy-to-implement therapies.

In this study, we demonstrated that post-stroke therapy with SCFAs have potential applications in stroke recovery. A previous study found that intestinal levels of acetate and propionate were significantly decreased in early phases of ischaemic stroke^13^. Intragastrically-administered butyrate for 14-days after stroke significantly improved stroke-induced brain damage and neurological deficits in cerebral ischaemic rats^13^. Moreover, treatment of endothelin-1-induced stroke rats by intraperitoneal injection of sodium butyrate at 6-hours post-stroke conferred neuroprotection and attenuated the loss of sensorimotor function^23^. Another study also demonstrated that bacteriotherapy of SCFA-producers in post-stroke aged mice reversed poor stroke recovery and reduced post-stroke neurological deficits and inflammation^24^. A key challenge for translation of these findings to patients is the delivery of SCFAs post-stroke. To achieve this, we tested the use of a HAMSAB diet, which we recently showed to reduce systolic blood pressure over a 3-week period in a phase II clinical trial in untreated hypertension patients^15^. Our results suggest that HAMSAB may represent a new opportunity for stroke recovery by both improving the gut epithelial barrier, infarct volume and neurogenesis, and reducing brain inflammation. Future trials will need to confirm their safety and potential benefits for stroke patients.

The blood-brain barrier (BBB) serves as a crucial physical barrier in maintaining homeostasis of the central nervous system. However, BBB is significantly damaged after a stroke^25^, which can exacerbate stroke-induced injury. In this study, we demonstrated that supplementation with SCFAs likely promotes recovery of BBB integrity, thereby attenuating stroke-induced BBB disruption and brain oedema. Previous studies found that treatment with butyrate, a histone deacetylase (HDAC) inhibitor, reduced stroke-induced BBB disruption^26^ and improved neurological outcomes^23^. Butyrate may act by reducing the expression of matrix metalloproteinase-9 (an enzyme that degrades extracellular matrix proteins), reducing the degradation of tight junction proteins and inhibiting NF-kB activation, similar to the effects of another HDAC inhibitor, valproic acid^26^.

Similarly, sodium butyrate-treated germ-free mice with hyperpermeable BBB showed increased occludin expression in the brain^27^. Moreover, propionate was found to be protective against lipopolysaccharide-induced disruption in BBB hCMEC/D3 cell culture^28^, supporting the idea that SCFAs regulate BBB integrity. These findings may have implications for other neurological diseases where the BBB is disrupted.

SCFAs regulate immune cells and are immune modulators to maintain homeostasis, which can impact both systemic inflammation and neuroimmunity^29^. A study showed that mixing the three major SCFAs (acetate, butyrate and propionate) promoted microglial maturation and restored defective microglia^30^. In particular, acetate not only promoted microglial maturation and regulated metabolic homeostatic state, but also regulated microglial phagocytosis and disease progression during neurodegeneration^31^. The impact of SCFAs on microglia activation might be closely linked to circulating lymphocytes and T cells recruitment^22^. In multiple sclerosis, propionate had protective effects by promoting the polarisation of naïve T cells toward anti-inflammatory T regulatory cells^32^. Although we did not investigate the role of microglia in our study, our results showed that supplementation with SCFAs significantly reduced neutrophil infiltration post-stroke. Considering neutrophil depletion reduces the extent of brain damage following traumatic brain injury^33^, our findings suggest that SCFAs may reduce brain damage and inflammation post-stroke.

There is limited knowledge about the importance of SCFA receptors in stroke recovery. A previous study found that GPR41 was highly expressed in the brain at 24-hours post-stroke, and that knockdown of GPR41 reversed the neuroprotective effect of sodium butyrate^34^. However, GPR41, GPR43 and GPR109A act on redundant signalling pathways, with co-expression in immune cells^16^. Thus, deleting just one or two of the receptors may mask their role in stroke. Due to the complex interplay of SCFAs and their receptors in regulating physiological processes^16^, we used a triple knockout mouse model, in which the three major SCFA receptors – GPR41, GPR43 and GPR109A – were removed to understand the importance of SCFA receptors in stroke recovery. We found that even on a high fibre diet, GPR41/43/109A^-/-^ mice had a worse stroke outcome, with increased brain inflammation and reduced neurogenesis. Loss of one or more receptors has been associated with dysregulated inflammatory responses to different immunologic challenges in inflammatory bowel disease^19, 35^ and elevated systolic blood pressure^36^. Mice lacking GPR41 receptors exhibit thickened aortas and increased collagen in blood vessels, leading to vascular fibrosis and high blood pressure^36^, which would increase the risk of an ischaemic stroke. Thus, our findings suggest that these receptors are essential in the fibre-gut microbiota-metabolite-stroke axis and may represent new targets for the development of drugs to aid in stroke recovery.

It is important to acknowledge that the present study has certain limitations. Firstly, contrary to previous studies in other disease models and patients using the same diets^15, 37^, we did not identify changes in the levels of plasma SCFAs in mice switched from a low fibre to a high fibre diet, a diet supplemented with SCFAs, or a diet containing HAMSAB. This could be because SCFAs are actively absorbed by the gut epithelial cells (which are more severely damaged after stroke compared to other diseases), liver and other systemic organs as an energy source^38^, and implies that the major beneficial effect of SCFA supplementation is associated with their action in the gut epithelia. Considering the amount of damage to the gut epithelial lining post-stroke and that the gut epithelial barrier was significantly improved with the administration of the various high fibre/SCFA-diets in this study, it is plausible that a large amount of SCFAs was directly used by the gut tissue. To determine this, future research should use isotope-labelled SCFAs to track their absorption and metabolism in stroke. Secondly, our study was only short-term (7-days post-stroke). Longer-term experiments are necessary to draw reliable conclusions about the benefits of gut microbial-produced metabolites in reversing poor stroke outcomes. Thirdly, our study only focused on young male mice. Future studies should investigate aged and female mice. This would provide valuable insights into the potential of diet intervention in post-stroke recovery and allow for a better understanding of the biological differences between males and females.

In conclusion, our research highlights the potential of high fibre dietary interventions, acting via the gut microbiota and production of SCFAs, to prevent and reverse the adverse effects of stroke. Moreover, we demonstrated the role of the SCFA receptors GPR41, GPR43 and GPR109A in stroke protection and recovery; these receptors could be potential targets for the development of novel stroke therapies. However, the downstream signalling pathways after activation of these receptors in the post-stroke brain remains unclear. Nonetheless, our study supports that SCFAs may be a new potent therapeutic option for stroke patients; this should be investigated in future clinical trials.

## Acknowledgements

We want to acknowledge the Monash Histology Facility for support with histology, the Monash Biomedical Imaging Facility for support with MRI imaging, Monash Proteomics and Metabolomics facility for support with SCFAs quantification, the Monash Animal Research Platform for support with animal work, and the Monash Bioinformatics Platform for access to M3 servers. The authors wish to acknowledge the use of the services and facilities of Australian Genome Research Facility the National Imaging Facility (NIF), a National Collaborative Research Infrastructure Strategy (NCRIS) capability at Monash University.

## Author contributions

A.P. planned and performed most of the in vivo and in vitro animal experiments and data analyses, provided intellectual inputs and wrote the manuscript. E.D. contributed to animal experimental work. H.J. and T.Z. contributed to 16S rRNA sequencing and analysis. D.A and D.J.C. contributed to SCFAs quantification. G.Z and M.dV. contributed to MRI imaging. C.R.M. contributed with knockout model and HAMSAB diet. B.R.S.B. and F.Z.M. conceived and supervised the research. B.K.H, B.R.S.B. and F.Z.M secured funding to support this study. All authors approved the final version of the manuscript.

## References

1. Feigin, V.L., et al. World Stroke Organization (WSO): Global Stroke Fact Sheet 2022. Int J Stroke 17, 18–29 (2022).

2. Peh, A., O’Donnell, J.A., Broughton, B.R.S. & Marques, F.Z. Gut Microbiota and Their Metabolites in Stroke: A Double-Edged Sword. Stroke 53, 1788–1801 (2022).

3. Singh, V., et al. Microbiota Dysbiosis Controls the Neuroinflammatory Response after Stroke. J Neurosci 36, 7428–7440 (2016).

4. Houlden, A., et al. Brain injury induces specific changes in the caecal microbiota of mice via altered autonomic activity and mucoprotein production. Brain Behav Immun 57, 10–20 (2016).

5. Jandhyala, S.M., et al. Role of the normal gut microbiota. World J Gastroenterol 21, 8787–8803 (2015).

6. Xie, L., Alam, M.J., Marques, F.Z. & Mackay, C.R. A major mechanism for immunomodulation: Dietary fibres and acid metabolites. Semin Immunol 66, 101737 (2023).

7. Peng, L., Li, Z.R., Green, R.S., Holzman, I.R. & Lin, J. Butyrate enhances the intestinal barrier by facilitating tight junction assembly via activation of AMP-activated protein kinase in Caco-2 cell monolayers. J Nutr 139, 1619–1625 (2009).

8. Wang, H.B., Wang, P.Y., Wang, X., Wan, Y.L. & Liu, Y.C. Butyrate enhances intestinal epithelial barrier function via up-regulation of tight junction protein Claudin-1 transcription. Dig Dis Sci 57, 3126–3135 (2012).

9. Desai, M.S., et al. A Dietary Fiber-Deprived Gut Microbiota Degrades the Colonic Mucus Barrier and Enhances Pathogen Susceptibility. Cell 167, 1339–1353 e1321 (2016).

10. Sonnenburg, E.D., et al. Diet-induced extinctions in the gut microbiota compound over generations. Nature 529, 212–215 (2016).

11. Ma, W., et al. Dietary fiber intake, the gut microbiome, and chronic systemic inflammation in a cohort of adult men. Genome Medicine 13, 102 (2021).

12. Reynolds, A., et al. Carbohydrate quality and human health: a series of systematic reviews and meta-analyses. The Lancet 393, 434–445 (2019).

13. Chen, R., et al. Transplantation of fecal microbiota rich in short chain fatty acids and butyric acid treat cerebral ischemic stroke by regulating gut microbiota. Pharmacol Res 148, 104403 (2019).

14. Threapleton, D.E., et al. Dietary fiber intake and risk of first stroke: a systematic review and meta-analysis. Stroke 44, 1360–1368 (2013).

15. Jama, H.A., et al. Prebiotic intervention with HAMSAB in untreated essential hypertensive patients assessed in a phase II randomized trial. Nature Cardiovascular Research 2, 35–43 (2023).

16. R. Muralitharan, R. & Marques, F.Z. Diet-related gut microbial metabolites and sensing in hypertension. Journal of Human Hypertension 35, 162–169 (2021).

17. Lewis, K., et al. Enhanced translocation of bacteria across metabolically stressed epithelia is reduced by butyrate†. Inflammatory Bowel Diseases 16, 1138–1148 (2009).

18. Silva, Y.P., Bernardi, A. & Frozza, R.L. The Role of Short-Chain Fatty Acids From Gut Microbiota in Gut-Brain Communication. Front Endocrinol (Lausanne) 11, 25 (2020).

19. Macia, L., et al. Metabolite-sensing receptors GPR43 and GPR109A facilitate dietary fibre-induced gut homeostasis through regulation of the inflammasome. Nat Commun 6, 6734 (2015).

20. Maslowski, K.M., et al. Regulation of inflammatory responses by gut microbiota and chemoattractant receptor GPR43. Nature 461, 1282–1286 (2009).

21. Suzuki, T., Yoshida, S. & Hara, H. Physiological concentrations of short-chain fatty acids immediately suppress colonic epithelial permeability. British Journal of Nutrition 100, 297–305 (2008).

22. Sadler, R., et al. Short-Chain Fatty Acids Improve Poststroke Recovery via Immunological Mechanisms. The Journal of Neuroscience 40, 1162–1173 (2020).

23. Park, M.J. & Sohrabji, F. The histone deacetylase inhibitor, sodium butyrate, exhibits neuroprotective effects for ischemic stroke in middle-aged female rats. Journal of Neuroinflammation 13, 300 (2016).

24. Lee, J., et al. Gut Microbiota-Derived Short-Chain Fatty Acids Promote Poststroke Recovery in Aged Mice. Circ Res 127, 453–465 (2020).

25. Peh, A., et al. Gut bacteria translocation to the brain after ischaemic stroke occurs via the sympathetic nervous system. bioRxiv, 2023.2004.2003.535309 (2023).

26. Wang, Z., Leng, Y., Tsai, L.K., Leeds, P. & Chuang, D.M. Valproic acid attenuates blood-brain barrier disruption in a rat model of transient focal cerebral ischemia: the roles of HDAC and MMP-9 inhibition. J Cereb Blood Flow Metab 31, 52–57 (2011).

27. Braniste, V., et al. The gut microbiota influences blood-brain barrier permeability in mice. Sci Transl Med 6, 263ra158 (2014).

28. Hoyles, L., et al. Microbiome-host systems interactions: protective effects of propionate upon the blood-brain barrier. Microbiome 6, 55 (2018).

29. Corrêa-Oliveira, R., Fachi, J.L., Vieira, A., Sato, F.T. & Vinolo, M.A. Regulation of immune cell function by short-chain fatty acids. Clin Transl Immunology 5, e73 (2016).

30. Erny, D., et al. Host microbiota constantly control maturation and function of microglia in the CNS. Nature Neuroscience 18, 965–977 (2015).

31. Erny, D., et al. Microbiota-derived acetate enables the metabolic fitness of the brain innate immune system during health and disease. Cell Metab 33, 2260–2276.e2267 (2021).

32. Haghikia, A., et al. Dietary Fatty Acids Directly Impact Central Nervous System Autoimmunity via the Small Intestine. Immunity 43, 817–829 (2015).

33. Kenne, E., Erlandsson, A., Lindbom, L., Hillered, L. & Clausen, F. Neutrophil depletion reduces edema formation and tissue loss following traumatic brain injury in mice. Journal of Neuroinflammation 9, 17 (2012).

34. Zhou, Z., et al. Sodium butyrate attenuated neuronal apoptosis via GPR41/Gβγ/PI3K/Akt pathway after MCAO in rats. J Cereb Blood Flow Metab 41, 267–281 (2021).

35. Kim, M.H., Kang, S.G., Park, J.H., Yanagisawa, M. & Kim, C.H. Short-chain fatty acids activate GPR41 and GPR43 on intestinal epithelial cells to promote inflammatory responses in mice. Gastroenterology 145, 396–406.e391-310 (2013).

36. Natarajan, N., et al. Microbial short chain fatty acid metabolites lower blood pressure via endothelial G protein-coupled receptor 41. Physiol Genomics 48, 826–834 (2016).

37. Thorburn, A.N., et al. Evidence that asthma is a developmental origin disease influenced by maternal diet and bacterial metabolites. Nature Communications 6, 7320 (2015).

38. Perry, R.J., et al. Acetate mediates a microbiome-brain-β-cell axis to promote metabolic syndrome. Nature 534, 213–217 (2016).

