## Supplementary material for "Dietary fibre reverses adverse post-stroke outcomes in mice via short-chain fatty acids and its sensing receptors GPR41, GPR43 and GPR109A": Online supplementary file

**Short title:** Dietary fibre in post-stroke recovery

Alex Peh^1,2^, Evany Dinakis^1^, Hamdi Jama^1^, Dovile Anderson^3^, Darren J. Creek^3^, Gang Zheng^4^, Michael de Veer^4^; Charles R. Mackay^5,6^, Tenghao Zheng^1^, Barbara Kemp-Harper^2^, Brad R.S. Broughton^2*^, Francine Z. Marques^1,7,8*^

^1^Hypertension Research Laboratory, School of Biological Sciences, Monash University, Melbourne, Australia; ^2^Cardiovascular & Pulmonary Pharmacology Group, Department of Pharmacology, Monash University, Melbourne, Australia; ^3^Monash Institute of Pharmaceutical Sciences, Monash University, Melbourne, Australia; ^4^Monash Biomedical Imaging, Monash University, Clayton, Victoria, Australia; ^5^Department of Microbiology, Biomedical Discovery Institute, Faculty of Medicine, Nursing and Health, Monash University, Clayton, Victoria, Australia; ^6^School of Pharmaceutical Sciences, Shandong Analysis and Test Center, Qilu University of Technology (Shandong Academy of Sciences), Jinan, 250014, China; ^7^Heart Failure Research Group, Baker Heart and Diabetes Institute, Melbourne, Australia; ^8^Victorian Heart Institute, Monash University, Melbourne, Australia.

*contributed equally as senior authors

**Correspondence to**: A/Prof Francine Marques, Hypertension Research Laboratory, School of Biological Sciences, Faculty of Science, Monash University, Melbourne, Australia, Phone: +61-03-9905 6958. E:

**Materials and methods**

**Animals**

All the experimental procedures performed in this study were approved by the Monash Animal Ethics Committee (animal ethics approval number: 23064) and followed the Australian Code for the Care and Use of Animals for Scientific Purposes. Wild-type (WT) C57BL6/J mice were purchased from the Monash Animal Research Platform (MARP, Melbourne, Australia). A triple knockout (GPR41/43/109A^-/-^) model that was bred on the C57BL6/J background, in which the three primary short-chain fatty acid (SCFA) receptors GPR41, GPR43, and GPR109A had been removed using CRISPR/Cas9 technology^1^, was obtained from our own specific pathogen-free breeding colony at Monash Animal Services. Male mice aged 8-9 weeks were used in all experiments and housed in an animal facility within Monash University.

**Diet treatment**

Experimental diets were ordered from Specialty Feeds, Australia. The nutrient composition of each diet is outlined in Extended Data Table 1. To demonstrate the beneficial effect of dietary fibre in stroke prevention and post-stroke recovery, the mice were fed with control (AIN93G) or nutrient-matched diets containing high fibre (HF) or low fibre (LF) for four weeks prior to surgery and fed with the same diet for seven days after the surgery. To determine whether dietary fibre can reverse adverse outcomes after stroke, the mice were first given low fibre before the surgery to achieve a poor post-stroke outcome and then shifted to either an HF diet, LF diet with SCFAs supplementation in drinking water (100mM sodium acetate and 100mM of sodium butyrate) or SCFAs-enriched high amylose maize starch - acetylated and butyrylated (HAMSAB) for seven days. HAMSAB diet was produced for this study specifically, on the background of the control (AIN93G) diet, with addition of 15% HAMSA and 15% HAMSB. The mice were assigned to different diets using a simple randomization method, and the specific combinations of diets are listed in Extended Data Table 2. Although the distinct colour and smell of the given diets made it impossible for researchers to be blinded to the diet treatments, steps were taken to minimize bias in downstream analysis by assigning each mouse a unique ID.

**Photothrombotic (PT) stroke mice model**

After receiving diet treatment for 28 days, mice were assigned to either a sham group or a stroke group using a simple randomization method. PT stroke was performed as previously described^2^. Briefly, the mice were anesthetised with isoflurane before any surgery. After removing the scalp and exposing the brain, we injected rose Bengal dye (10mg/µl/g body weight) intraperitoneally. To activate the dye, we focused illumination on an area of 1.5mm diameter in the M1 cortex on the intact skull for 15 minutes via a 20x objective to activate the dye. The mice were placed on a heating pad throughout the procedure to maintain their body temperature at 37^o^C. For sham-operated surgery, the same procedure was performed except for no illumination. Mice were only excluded from the analysis if they died before the 7-day study endpoint. While all the sham mice survived, we have several mortalities in the following groups: CTL-CTL (5 dead mice); LF-LF (5 dead mice); HF-HF (3 dead mice); LF-HF (2 dead mice); LF-HAMSAB (1 dead mouse); GPR41/43/109A^-/-^ (2 dead mice).

**Motor function assessment**

To assess the post-surgery improvement in motor function of the mice, we performed the wire-hanging test at day 0 (D0; baseline), D1, D3 and D7. The wire-hanging test was performed as previously described^2^ with slight modification. In brief, the mice were hung on a wire stretched between two posts approximately 60cm high. The mice were able to hang their weight on wires with their forelimbs. The time for each mouse to fall off the wire was recorded for up to a maximum of 300sec. Three trials were performed for each mouse with a 5 mins resting interval between trials, and their average hanging time was calculated for analysis.

**Magnetic resonance imaging (MRI)**

The brain infarct developed at D1 post-stroke was determined by MRI imaging. All MRI experiments were performed on a Bruker 9.4T animal MRI Scanner (Bruker BioSpin GmbH, Ettlingen, Germany) equipped with a Bruker 86mm volumetric transmitter coil and a single-channel mouse head receiver coil. All mice were maintained under 1-1.5% isoflurane anaesthesia throughout the imaging procedure. We monitored the respiration and body temperature of the mice throughout the experiment. The Bruker 2D T2-weighted RARE imaging sequence was used with the following parameters: TE/TR = 36 ms/7000 ms; RARE factor= 8; FOV= 16.8 x 16.8 mm^2^; number of slices = 26; slice thickness= 0.5mm; matrix= 168 x 168; averages = 4; orientation = Axial. We analysed the MRI data using MRIcron software to determine the brain infarct size and brain oedema.

**Infarct volume and brain oedema**

To determine the brain infarct size and brain oedema at D7, frozen brains were cut into 30 μm coronal sections on a cryostat. Evenly spaced sections (separated by 210 µm) spanning the infarct were thaw-mounted onto Superfront-OT plus microscope slides and stained with 0.1% thionin. Images were captured and analysed using ImageJ software (version 1.53t; National Institutes of Health). Brain oedema was calculated by comparing the ischaemic and contralateral hemispheres using the formula: Brain oedema (as % to contralateral hemisphere) = [(volume of ischaemic hemisphere – volume of contralateral hemisphere)/ volume of contralateral hemisphere] * 100%. % infarct recovery = (different in infarct size between D1 and D7 / infarct size at D1) * 100%. A similar formula was used to calculate the recovery rate for brain oedema.

**Tissue collection**

All samples were collected after the mice were euthanised with an overdose of isoflurane and perfused with clean phosphate-buffered saline (PBS). The weight or length of the organs were measured and recorded. Samples were either snap-frozen in liquid nitrogen and stored at -80^o^C until further analysis, fixed in 10% formalin before paraffin embedding, or direct OCT embedding for histopathology studies. Blood plasma was collected after centrifugation at 4^o^C, 1500 rpm for 15 minutes, and stored at -80^o^C before being sent for SCFA quantification.

**Immunofluorescence labelling**

All immunofluorescence labelling was performed after optimisation according to the manufacturer’s protocols. Briefly, three to four 10µm brain tissue sections collected from the middle of the infarct area were first air-dried before being fixed with 4% paraformaldehyde for 15 minutes. The sections were washed with 0.01M PBS (3x 10 minutes) and blocked with 10% goat serum (Abcam; Cambridge, USA, diluted in 0.01M PBS) for 1 hour. The sections were then incubated with the desired primary antibody overnight at room temperature in a humidified box (see Extended Data Table 3). The following day, sections were washed in 0.01M PBS (3x 10 minutes) and incubated with the appropriate secondary antibody at room temperature for 2 hours. The primary and secondary antibodies used are summarised in Extended Data Table 1. Sections were washed in 0.01M PBS (3x 10 minutes) and then carefully dried before being mounted in Vectashield Antifade Mounting Medium with DAPI (Vector Laboratories, Burlingame, CA). A similar procedure was employed to immunolabel the colon section. In brief, the tissue was air-dried and fixed in 10% formalin for 20 minutes. After washing, the tissue was labelled with a primary antibody (see Extended Data Table 3) overnight, followed by washing and labelling with a secondary antibody (see Extended Data Table 3) for 2 hours at room temperature before mounted in a mounting medium and cover-slipped. Negative controls were performed using the same protocol in the absence of primary antibody and showed no immunofluorescence (see Extended Data Fig. 11).

**Gut histology analysis**

The small intestine was separated into duodenum, jejunum and ileum, fixed in 10% formalin for 24 hrs, embedded in paraffin, sectioned at 10μm and stained for Periodic Acid-Schiff Alcian Blue (PAS/AB) and Masson’s trichrome. Whole tissue sections were scanned at 40x magnification with the Scanscope AT Turbo (Aperio). Each intestinal tissue section was captured at 10x magnification in three different fields of view using ImageScope software (Aperio, v12.4.0.5043). The number of goblet cells per millimetre square area, villi length, intestinal muscularis propria thickness, and fibrosis thickness were quantified manually using ImageJ software.

**Quantification of SCFAs**

SCFAs in the plasma were quantified by employing a derivatising agent and its ^13^C_6_-labelled version 3-NPH/^13^C_6_-3NPH as previously described^3^ using QExactive mass spectrometer (Thermo Fisher Scientific) with Dionex UltiMate 3000 RSLC nano system (Thermo Fisher Scientific) as recently described^4^. Briefly, 3-NPH was used to derivatize SCFAs in plasma, spiked plasma QC samples, and calibrators in water. At the same time, a mixture of SCFAs in water was derivatized with ^13^C_6_-3-NPH to generate an isotopically labelled internal standard mixture. An equal amount of this mixture was added to all samples, QCs and calibrators. Mass spectral data was collected using PRM (Parallel Reaction Monitoring) scan mode and quantified by integrating the diagnostic fragment ions generated from each precursor ion with expected retention time (137.0357 m/z for the target compound 143.0558 m/z for its internal standard with a mass error window of 10 ppm).

**Microbial DNA extraction and 16S sequencing**

Microbial DNA extraction of caecum samples collected at euthanasia was performed using a DNeasy PowerSoil DNA isolation kit (Qiagen) according to the protocol provided by the manufacturer. DNA contents of each sample were quantified using Nanodrop (ThermoFisher Scientific). The V4 region of the 16S ribosomal RNA was amplified with 515F and 806R primers and 10ng of the microbial DNA and Platinum™ Hot Start PCR Master Mix (ThermoFisher Scientific), according to the protocol of Earth Microbiome Project^5^. After the amplification, 240ng of the library was pooled and then cleaned up using QIAquick PCR Purification Kit (Qiagen) before being sequenced at the Australian Genomic Research Facility (AGRF, Melbourne, Australia) using an Illumina MiSeq sequencer (300-bp paired-end reads).

**16S rRNA bioinformatics analyses**

The microbiome sequencing data were analysed using QIIME2 (2020.2 version) in R Studio (version 1.2.1335)^6^. The raw reads were trimmed to have at least 20 Phred quality scores, reads were merged, and the chimeric reads were removed. DADA2 QIIME2 plug-in was used to remove and correct noisy reads^7^. Alpha‐diversity metrics (Faith's phylogenetic diversity index, Shannon diversity index, Observed species richness, and Species Evenness index), and beta-diversity metrics (both weighted UniFrac and unweighted UniFrac) shown as principle coordinate analysis (PCoA) plots, were estimated using the q2‐diversity QIIME2 plug-in after rarefying the samples to a minimum depth of 8,000 amplicon sequence variants (ASVs). MicrobiomeAnalyst online software^8^ was used to visualise taxonomic bar charts at both phylum and genus levels, as well as to perform a Linear discriminant analysis Effect Size (LEfSe) analysis with a threshold of 2.0 on the logarithmic LDA score for discriminative features and a p-value cut off of 0.05 and FDR-adjusted.

**Statistical analyses**

Quantitative data were presented as mean ± standard error of the mean (SEM). Statistical analyses were performed using GraphPad Prism version 9.1.0 software. Statistical comparisons were made either by two-tailed unpaired t-test (when comparing two groups), by one-way ANOVA corrected by false discovery rate (FDR, when comparing more than two groups; two-stage step-up method of Benjamini, Krieger and Yekutieli), or by two-way ANOVA corrected by FDR (when comparing two grouping variables; two-stage step-up method of Benjamini, Krieger and Yekutieli). The permutational multivariate analysis of variance (PERMANOVA) test was used for microbiome analysis. *P*<0.05 was considered statistically significant.

**Data availability**

All sequencing data was deposited in NCBI Sequence Read Archive database SUB13166781 (BioProject ID PRJNA961228). Other data are available from the corresponding author on request.

**Extended Data Tables**

**Extended Data Table 1. Nutrient composition of experimental diets.**

| **Nutritional Parameter / Diet** | **Control diet** | **High fibre** | **Low fibre** |
| --- | --- | --- | --- |
| Code | AIN93G | SF11-025 (AIN93G background) | SF09-028 (AIN93G background) |
| Sucrose | 10.0% | - | - |
| Casein (Acid) | 20.0% | 20.0% | 20.0% |
| Canola Oil | 7.0% | 7.0% | 7.0% |
| Cellulose | 5.0% | 5.0% | - |
| Starch | 40.4% (wheat, non-resistant starch) | 63.6% (gel crisp, 100% resistant starch) | - |
| Dextrinised Starch | 13.2% (non-resistant starch) | - | 68.6% (non-resistant starch) |
| L-Methionine | 0.3% | 0.3% | 0.3% |
| Trace Minerals | 0.1% | 0.1% | 0.1% |
| Fine Calcium Carbonate | 1.3% | 1.3% | 1.3% |
| Salt (Fine Sodium Chloride) | 0.3% | 0.3% | 0.3% |
| Potassium Dihydrogen Phosphate | 0.7% | 0.7% | 0.7% |
| Potassium Sulphate | 0.2% | 0.2% | 0.2% |
| Potassium Citrate | 0.2% | 0.2% | 0.2% |
| Vitamins | 1.0% | 1.0% | 1.0% |
| Choline Chloride 75% w/w | 0.3% | 0.3% | 0.3% |

**Extended Data Table 2. An overview of diet treatment combinations given.**

| **Group** | **Treatment 1**  **(28 days pre-surgery treatment)** | **Treatment 2**  **(7 days post-surgery treatment)** |
| --- | --- | --- |
| CTL-CTL | AIN93G | AIN93G |
| HF-HF | High fibre (HF) | High fibre (HF) |
| LF-LF | Low fibre (LF) | Low fibre (LF) |
| LF-HF | Low fibre (LF) | High fibre (HF) |
| LF-LF+SCFAs | Low fibre (LF) | Low fibre (LF)+ SCFAs |
| LF-HAMSAB | Low fibre (LF) | HAMSAB |

*Legend: CTL, control; HAMSAB, acetylated and butyrylated high amylose maize starch; HF, high fibre; LF, low fibre; SCFAs, short-chain fatty acids.

**Extended Data Table 3. List of primary and secondary antibodies used for immunofluorescence.**

| Tissue | Primary antibody | Dilution | Incubation temperature | Secondary antibody | Dilution |
| --- | --- | --- | --- | --- | --- |
| Brain | *Rabbit anti-MPO (Abcam; ab9535)* | 1:200 | Room temperature | Goat anti-rabbit IgG Alexa Fluor 594 (Life Technologies) | 1:500 |
|  | *Rabbit anti-DCX (Abcam; ab18723)* | 1:500 | 4˚C | Goat anti-rabbit IgG Alexa Fluor 594 (Life Technologies) | 1:500 |
|  | *Rabbit anti-mouse ZO-1 (61-7300)* | 1:100 | 4^o^C | Goat anti-rabbit IgG Alexa Fluor 594 (Life Technologies) | 1:500 |
| Colon | *Rabbit anti-mouse ZO-1 (61-7300)* | 1:100 | 4^o^C | Goat anti-rabbit IgG Alexa Fluor 594 (Life Technologies) | 1:500 |
|  | *Rat anti-mouse CD32/EpCam (14-5791-81)* | 1:500 | 4^o^C | Goat anti-rat IgG Alexa Fluor 488 (Life Technologies) | 1:500 |

*Legend: DCX, Doublecortin; EpCam, epithelial cellular adhesion molecule; MPO, myeloperoxidase; ZO-1, zonula occludens-1

**Extended Data Figures**

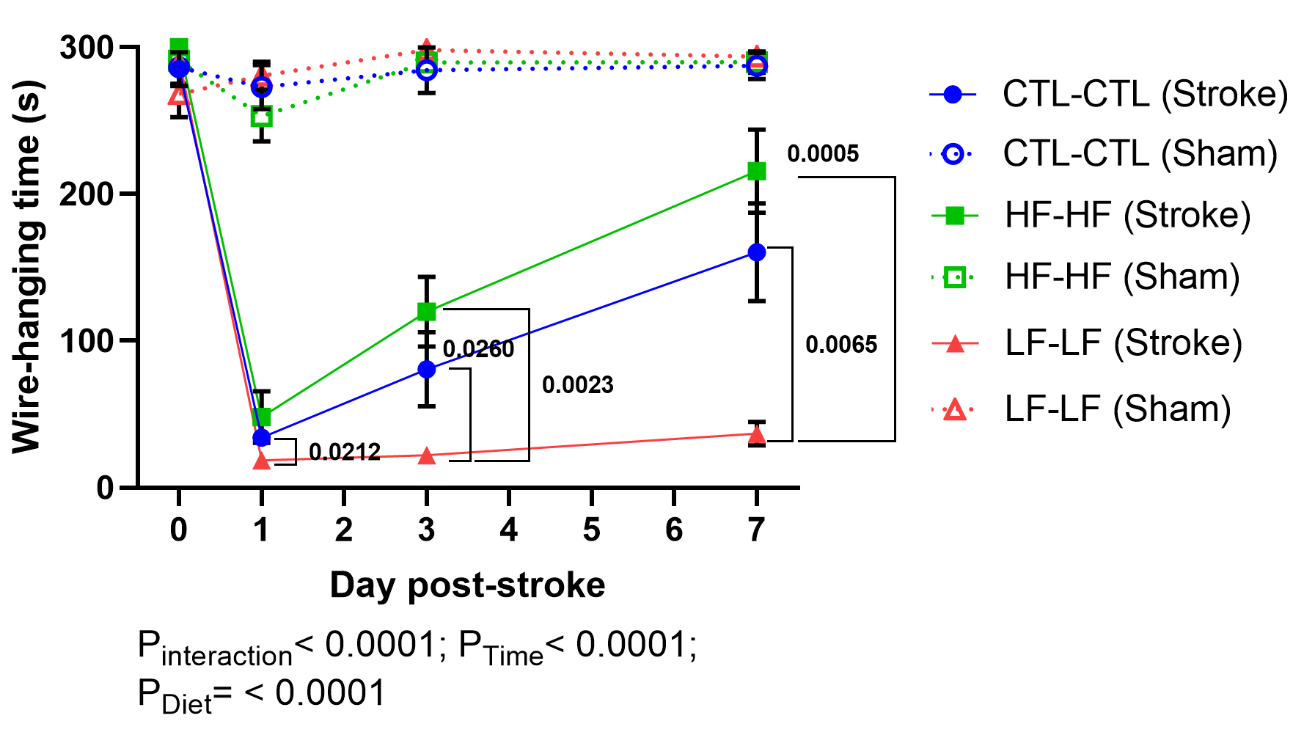

**Extended Data Fig. 1. Motor functional test of CTL-CTL, LF-LF, and HF-HF.**

The wire-hanging test was carried out after sham-operated surgery or PT stroke surgery at D0 (baseline), D1, D3 and D7. Sample size= 8/group; error bar denotes mean±SEM. Two-way ANOVA was used to analyse data. The graph displays only the significant values within the PT stroke group to prevent visual clutter. Legend: CTL, control; HF, high fibre; LF, low fibre.

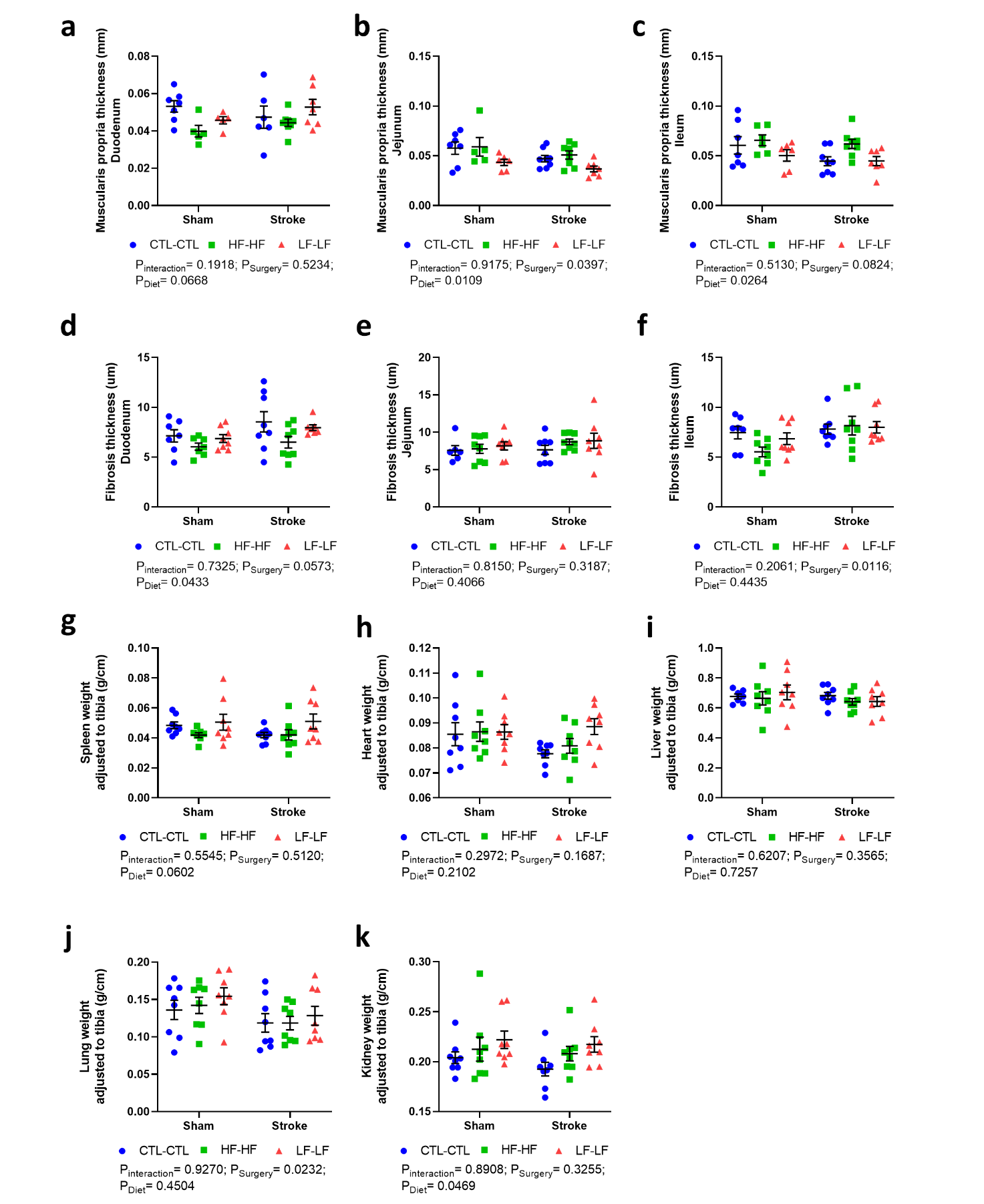

**Extended Data Fig. 2. Muscularis propria thickness, fibrosis thickness and organ weight between CTL-CTL, HF-HF and LF-LF groups.**

No difference in **a-c,** muscularis propria layer thickness, **d-f,** fibrosis thickness and various organs weight adjusted to tibia length: **g,** spleen, **h,** heart, **i,** liver, **j,** lung and **k,** kidney. Sample size= 8/group; error bar denotes mean±SEM. Two-way ANOVA was used to analyse data. Legend: CTL, control; HF, high fibre; LF, low fibre.

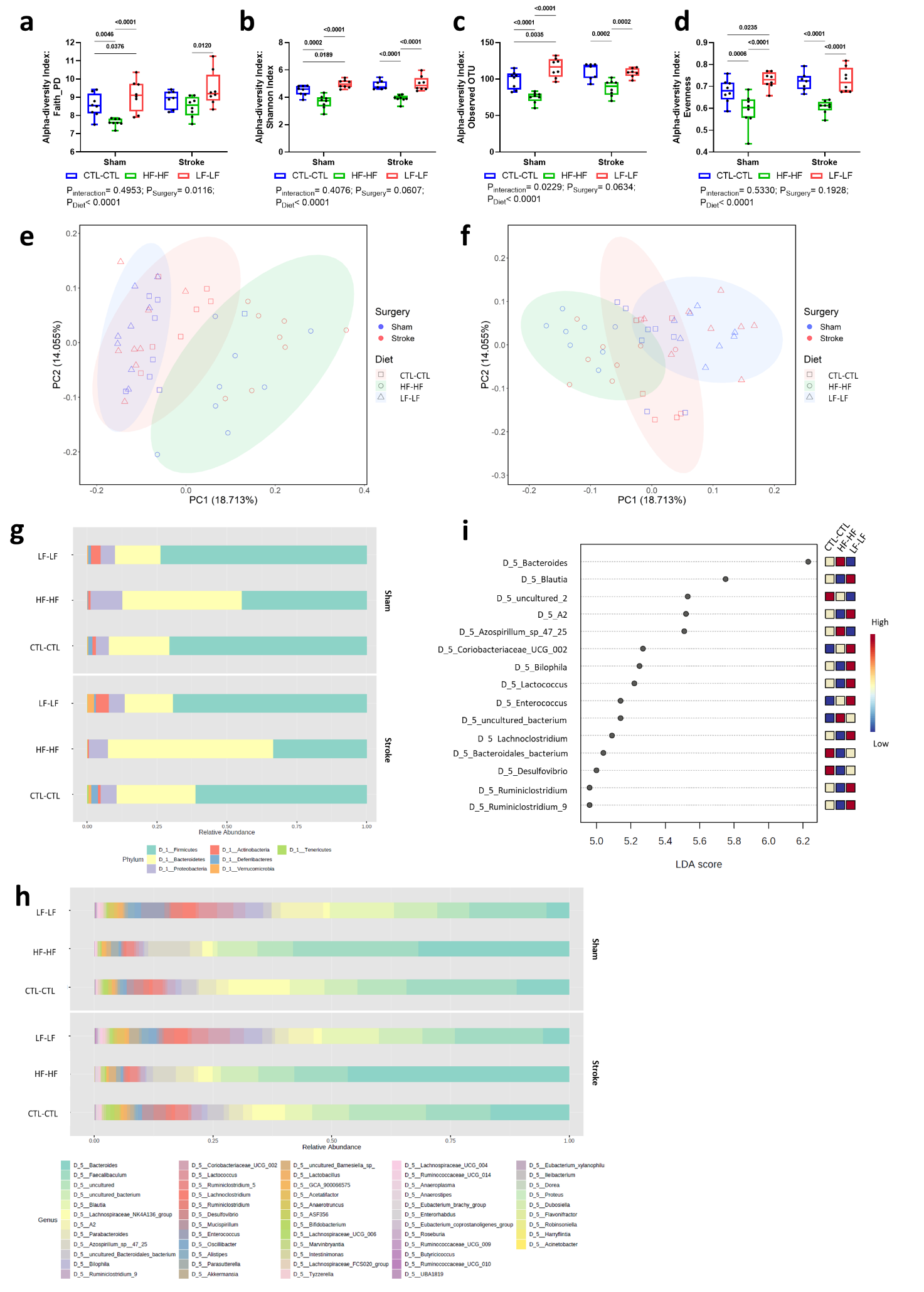

**Extended Data Fig. 3. α and β diversity profile of the CTL-CTL, HF-HF, and LF-LF gut microbiome at D7 post-surgery.**

Box plots illustrating α diversity of gut microbiome between CTL-CTL, HF-HF, and LF-LF: **a,** Faith's phylogenetic diversity; **b,** Shannon diversity index; **c,** Observed species richness; and **d,** Species evenness index. Sample size= 8/group; error bar denotes median and 95% CI. Two-way ANOVA was used to analyse data. PCoA plots showing the 95% CI β-diversity clustering patterns of samples according to the type of diet treatment, based on **e,** weighted UniFrac distance (p-value: 0.001); and **f,** unweighted UniFrac distance (p-value: 0.001). Taxa bar plot that is classified based on **g,** phylum-level and **h,** genus-level. **i,** Linear Discriminant Analysis (LDA) Effect Size (LEfSe) bar plot identified the top 15 significant features classified based on the diet treatment (threshold on the logarithmic LDA score for discriminative features: 2.0; p-value cutoff: 0.05, FDR-adjusted). Legend: CTL, control; HF, high fibre; LF, low fibre.

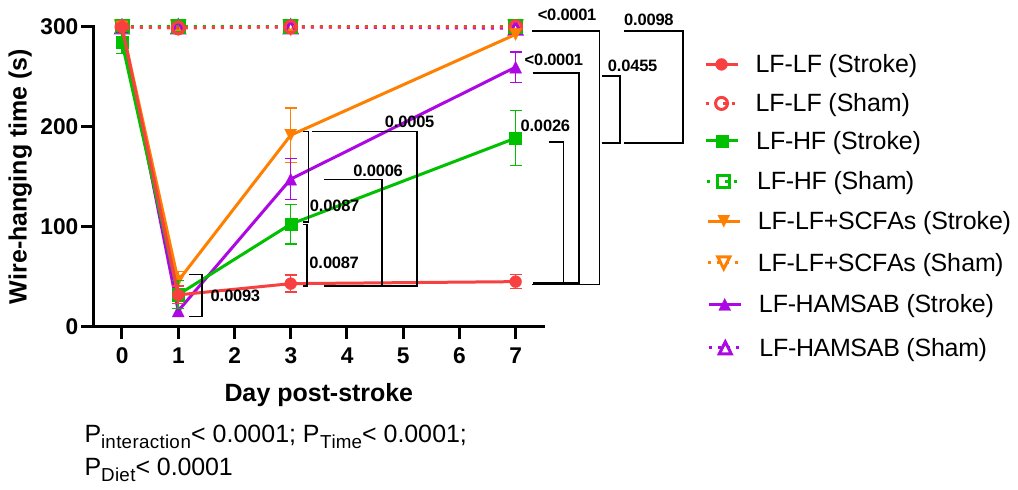

**Extended Data Fig. 4. Motor functional test of LF-LF, LF-HF, LF-LF+SCFAs and LF-HAMSAB.**

The wire-hanging test was carried out after sham-operated surgery or PT stroke surgery at D0 (baseline), D1, D3 and D7. Sample size= 8/group; error bar denotes mean±SEM. Two-way ANOVA was used to analyse data. The graph displays only the significant values within the PT stroke group to prevent visual clutter. Legend: LF, low fibre; HF, high fibre; SCFAs, short-chain fatty acids; HAMSAB, acetylated and butyrylated high amylose maize starch.

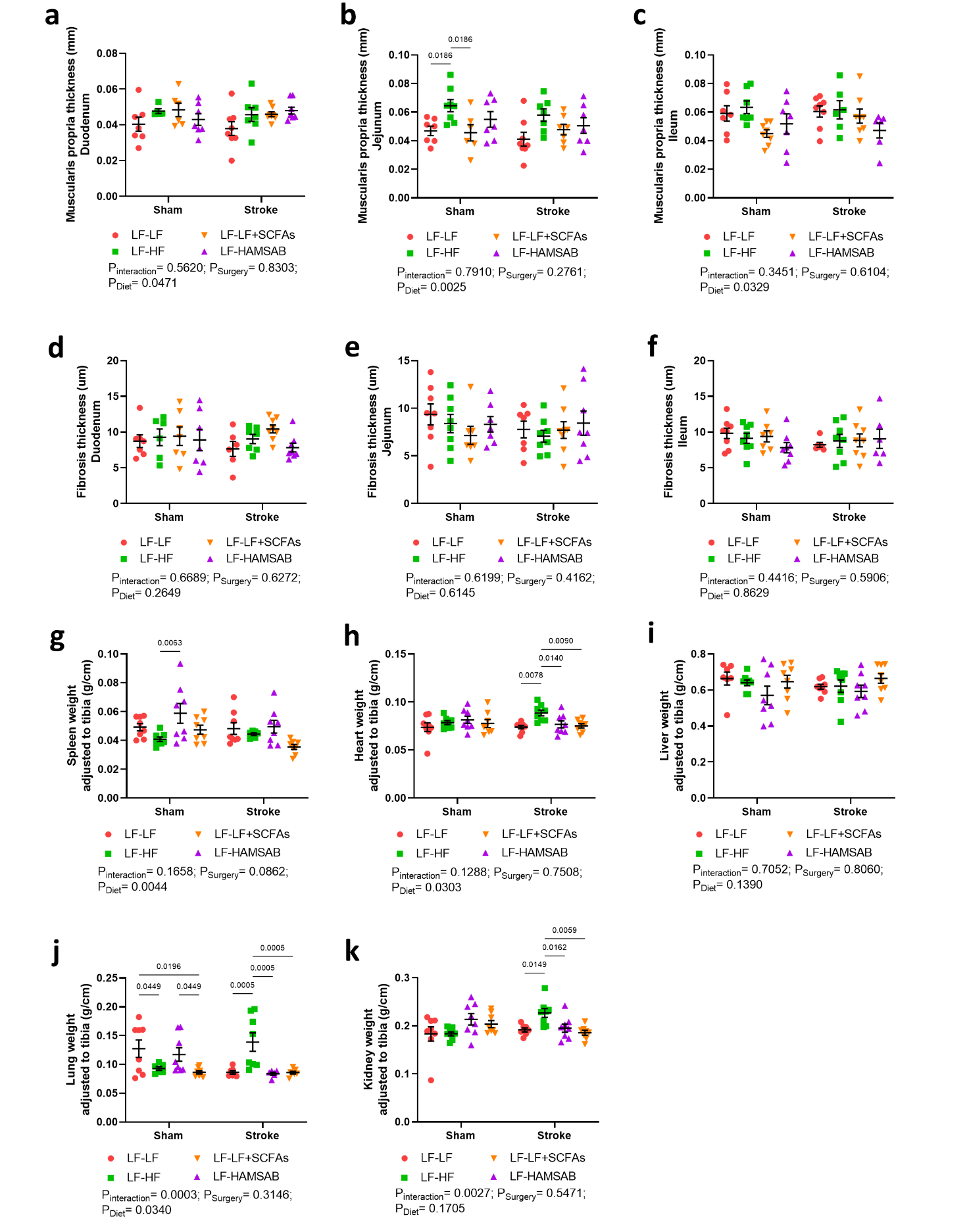

**Extended Data Fig. 5.** **Muscularis propria thickness, fibrosis thickness and organ weight between LF-LF, LF-HF, LF-LF+SCFAs and LF-HAMSAB groups.**

Quantification of **a-c,** muscularis propria layer thickness, **d-f,** fibrosis thickness and various organs weight adjusted to tibia length: **g,** spleen, **h,** heart, **i,** liver, **j,** lung and **k,** kidney. Sample size= 8/group; error bar denotes mean±SEM. Two-way ANOVA was used to analyse data. Legend: LF, low fibre; HF, high fibre; SCFAs, short-chain fatty acids; HAMSAB, acetylated and butyrylated high amylose maize starch.

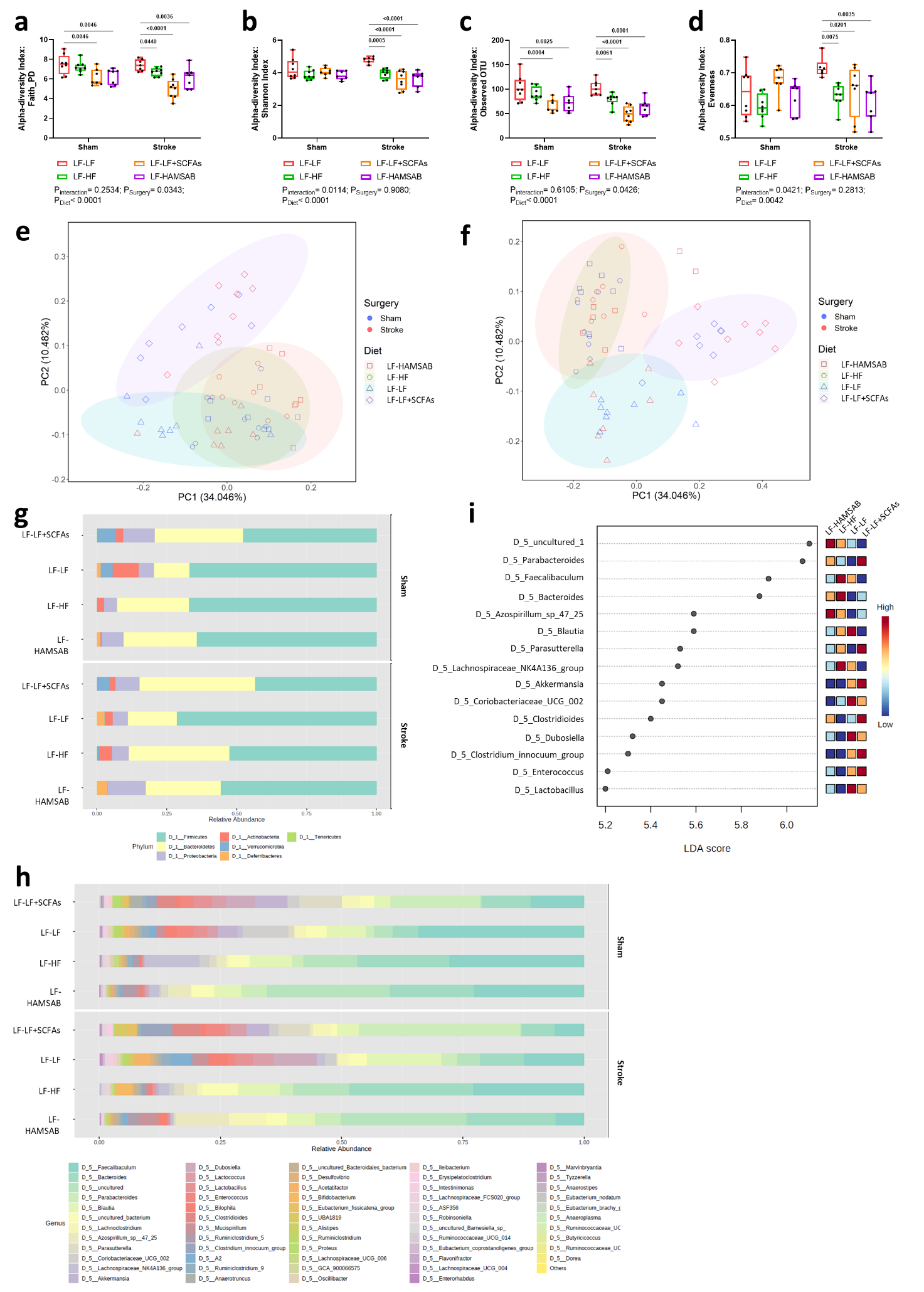

**Extended Data Fig. 6. α and β diversity profile of the gut microbiome of LF-LF, LF-HF, LF-LF+SCFAs and LF-HAMSAB at D7 post-surgery.**

Box plots illustrating α diversity of gut microbiome between LF-LF, LF-HF, LF-LF+SCFAs and LF-HAMSAB: **a,** Faith's phylogenetic diversity; **b,** Shannon diversity index; **c,** Observed species richness; and **d,** Species evenness index. Sample size= 8/group; error bar denotes median and 95% CI. Two-way ANOVA was used to analyse data. PCoA plots showing the 95% CI β-diversity clustering patterns of samples according to the type of diet treatment, based on **e,** weighted UniFrac distance (p-value: 0.001); and **f,** unweighted UniFrac distance (p-value: 0.001). Taxa bar plot that is classified based on **g,** phylum-level and **h,** genus-level. **i,** Linear Discriminant Analysis (LDA) Effect Size (LEfSe) bar plot identified the top 15 significant features classified based on the diet treatment (threshold on the logarithmic LDA score for discriminative features: 2.0; p-value cutoff: 0.05, FDR-adjusted). Legend: LF, low fibre; HF, high fibre; SCFAs, short-chain fatty acids; HAMSAB, acetylated and butyrylated high amylose maize starch.

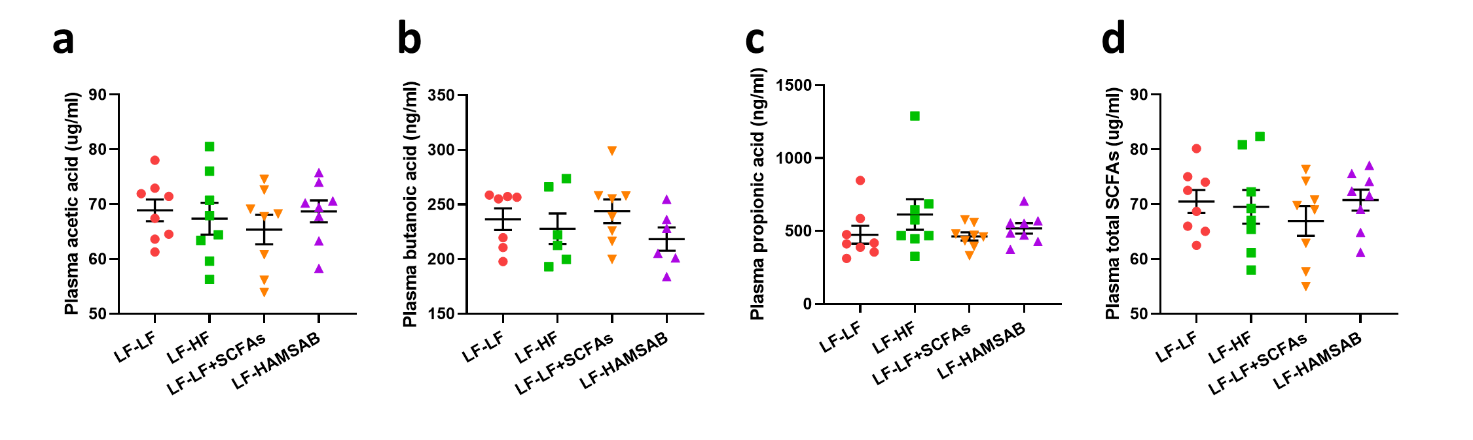

**Extended Data Fig. 7. No difference was detected in the SCFAs level from circulating blood plasma of LF-LF, LF-HF, LF-LF+SCFAs and LF-HAMSAB.**

Quantification of **a,** acetic acid, **b,** butanoic acid, **c,** propionic acid, and **d,** total SCFAs from the peripheral blood circulation. Legend: LF, low fibre; HF, high fibre; SCFAs, short-chain fatty acids; HAMSAB, acetylated and butyrylated high amylose maize starch.

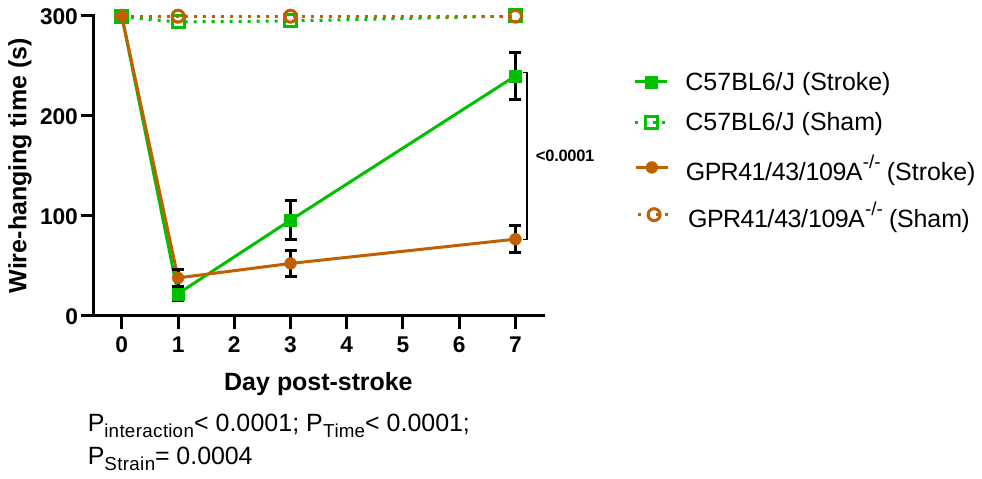
**Extended Data Fig. 8. Motor functional test of C57BL6/J and** **GPR41/43/109A^-/-^.**

The wire-hanging test was carried out after sham-operated surgery or PT stroke surgery at D0 (baseline), D1, D3 and D7. Sample size= 8/group; error bar denotes mean±SEM. Two-way ANOVA was used to analyse data. The graph displays only the significant values within the PT stroke group to prevent visual clutter.

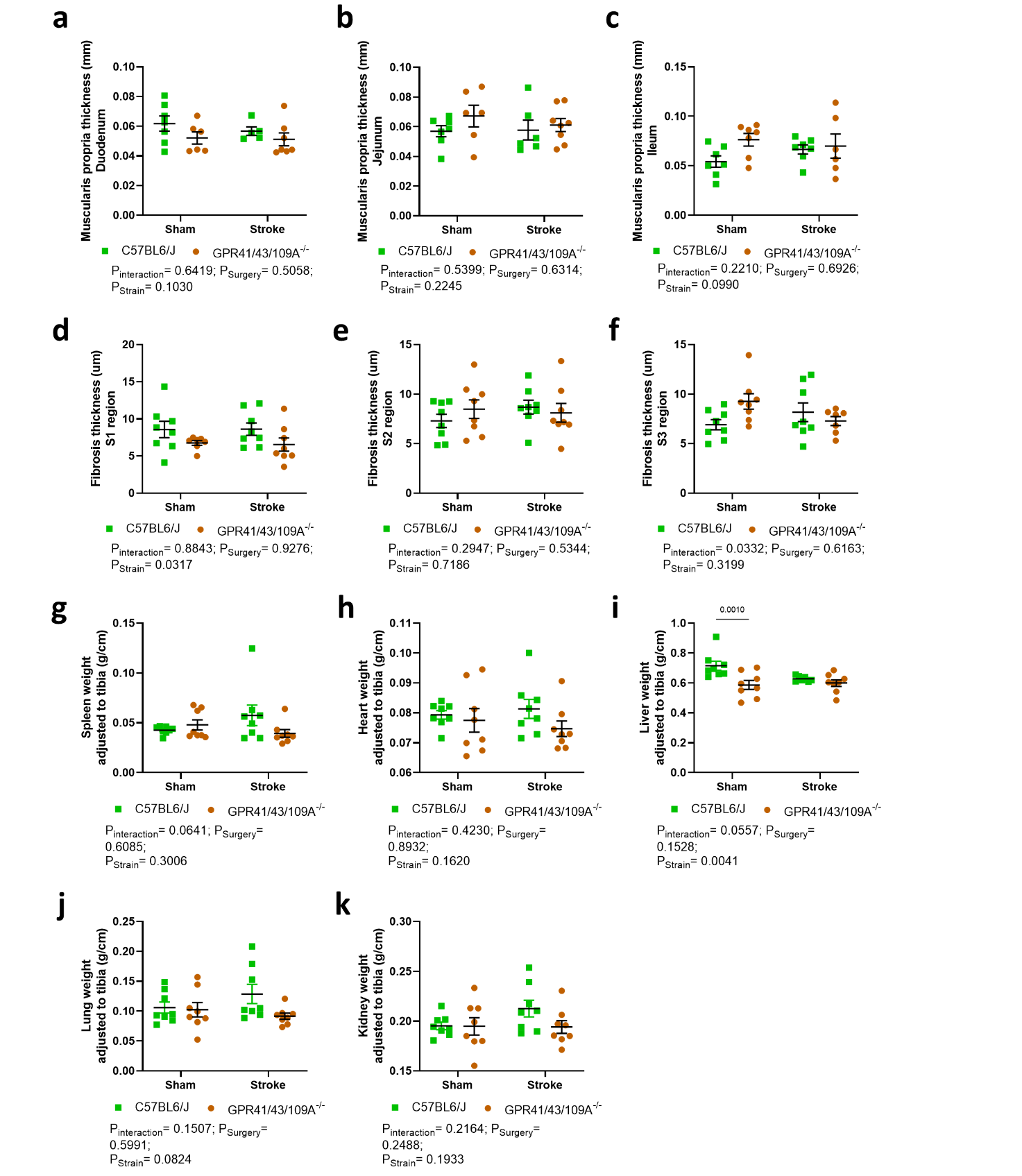

**Extended Data Fig. 9.** **Muscularis propria thickness, fibrosis thickness and organ weight between C57BL6/J and GPR41/43/109A^-/-^ groups.**

Quantification of **a-c,** muscularis propria layer thickness, **d-f,** fibrosis thickness and various organs weight adjusted to tibia length: **g,** spleen, **h,** heart, **i,** liver, **j,** lung and **k,** kidney. Sample size= 8/group; error bar denotes mean±SEM. Two-way ANOVA was used to analyse data.

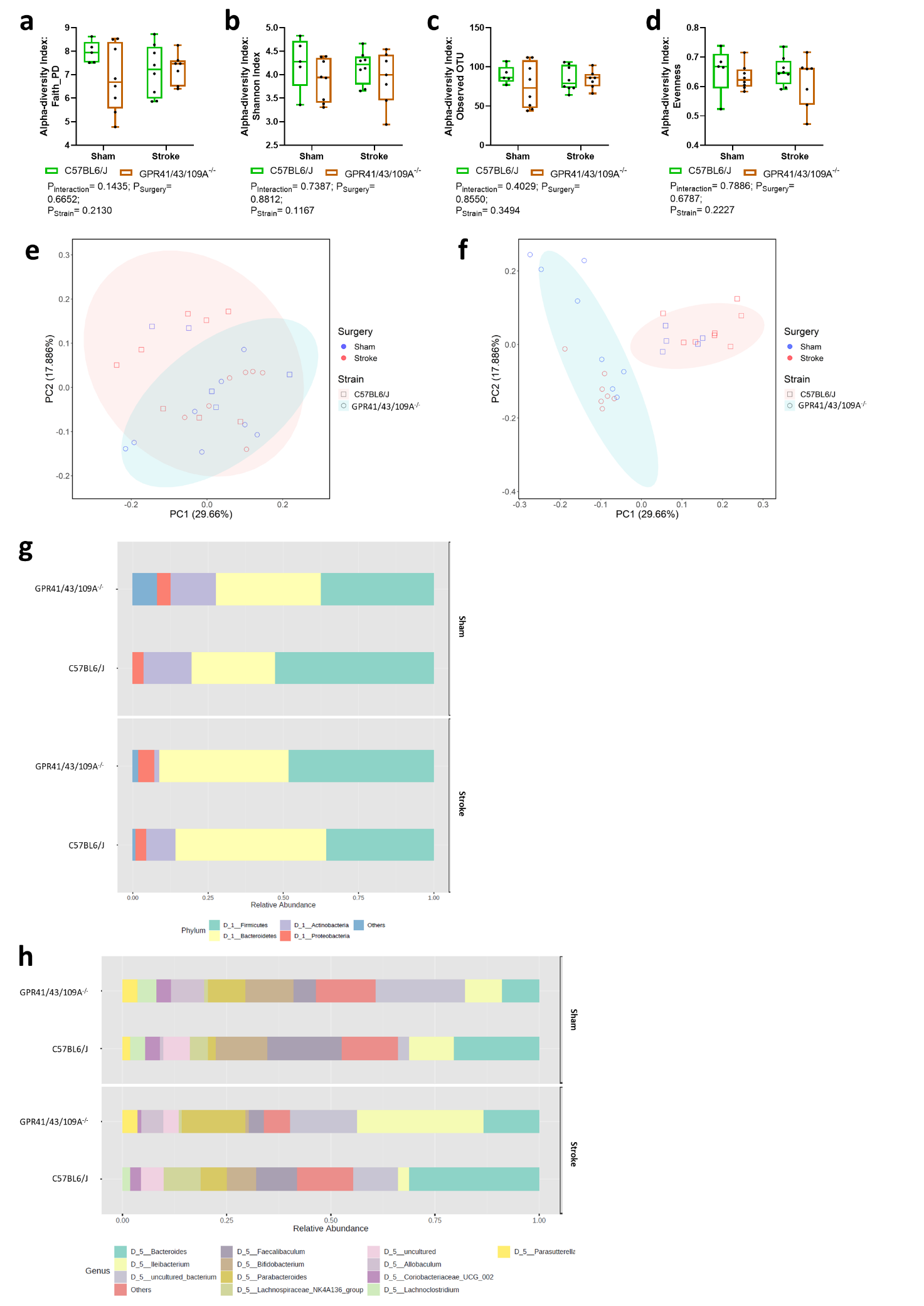

**Extended Data Fig. 10. α and β diversity profile of the gut microbiome in C57BL6/J and GPR41/43/109A^-/-^ mice at D7 post-surgery.**

Box plots illustrating α diversity of gut microbiome between C57BL6/J and GPR41/43/109A^-/-^ mice: **a,** Faith's phylogenetic diversity; **b,** Shannon diversity index; **c,** Observed species richness; and **d,** Species evenness index. Sample size= 8/group; error bar denotes median and 95% CI. Two-way ANOVA was used to analyse data. PCoA plots showing the 95% CI β-diversity clustering patterns of samples according to the strain, based on **e,** weighted UniFrac distance (p-value: 0.003); and **f,** unweighted UniFrac distance (p-value: 0.001). Taxa bar plot that is classified based on **g,** phylum-level and **h,** genus-level. No significant features were identified in LefSe analysis.

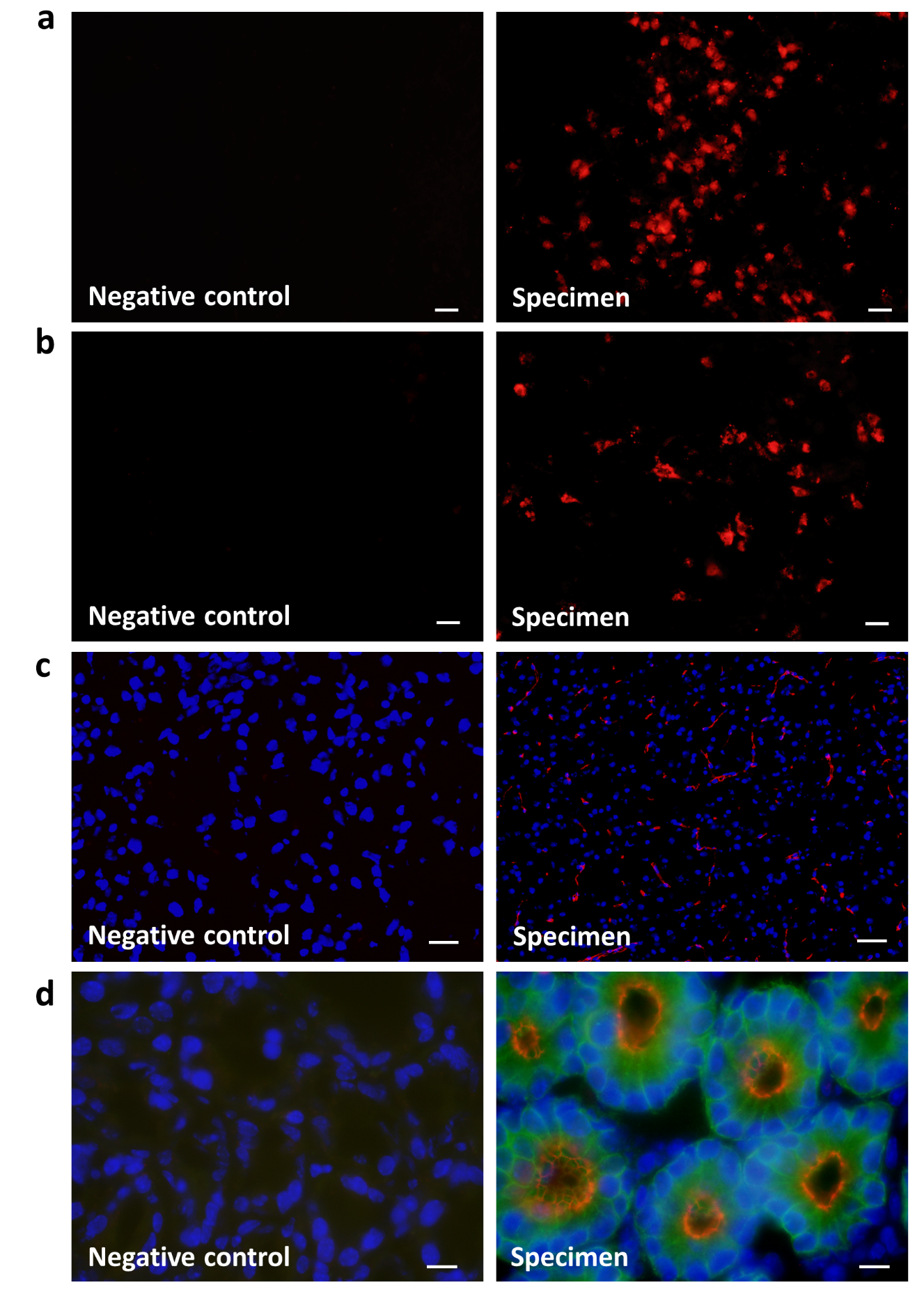

**Supplementary Fig. 11. Negative controls for immunofluorescence labelling.**

The results of negative controls indicated a lack of immunofluorescence for the following markers in the brain: **a,** MPO, **b,** DCX, and **c,** ZO-1, as well as in the colon: **d,** ZO-1 and EpCam.

**References**

1.	Maslowski, K.M.*, et al.* Regulation of inflammatory responses by gut microbiota and chemoattractant receptor GPR43. *Nature* **461**, 1282-1286 (2009).

2.	Peh, A.*, et al.* Gut bacteria translocation to the brain after ischaemic stroke occurs via the sympathetic nervous system. *bioRxiv*, 2023.2004.2003.535309 (2023).

3.	Han, J., Lin, K., Sequeira, C. & Borchers, C.H. An isotope-labeled chemical derivatization method for the quantitation of short-chain fatty acids in human feces by liquid chromatography-tandem mass spectrometry. *Anal Chim Acta* **854**, 86-94 (2015).

4.	Jama, H.*, et al.* Maternal diet and gut microbiota influence predisposition to cardiovascular disease in the offspring. *bioRxiv*, 2022.2003.2012.480450 (2022).

5.	Caporaso, J.G.*, et al.* Global patterns of 16S rRNA diversity at a depth of millions of sequences per sample. *Proc Natl Acad Sci U S A* **108 Suppl 1**, 4516-4522 (2011).

6.	Bolyen, E.*, et al.* Reproducible, interactive, scalable and extensible microbiome data science using QIIME 2. *Nature Biotechnology* **37**, 852-857 (2019).

7.	Callahan, B.J.*, et al.* DADA2: High-resolution sample inference from Illumina amplicon data. *Nat Methods* **13**, 581-583 (2016).

8.	Chong, J., Liu, P., Zhou, G. & Xia, J. Using MicrobiomeAnalyst for comprehensive statistical, functional, and meta-analysis of microbiome data. *Nature Protocols* **15**, 799-821 (2020).
